# A spatially-resolved blueprint of the developing human lung reveals a WNT-driven niche for basal stem cells

**DOI:** 10.1101/2024.10.01.612096

**Authors:** Peggy P. Hsu, Ansley S. Conchola, Tristan Frum, Xiangning Dong, Lila Tudrick, Varun Ponnusamy, Michael S. Downey, Manqi Wu, Mengkun Yang, Yusoo Lee, Emma Niestroy, Yu-Hwai Tsai, Angeline Wu, Sha Huang, Ian A. Glass, Sofia D. Merajver, Jason R. Spence

## Abstract

Organs are composed of diverse cell types that change across space and time during development. To interrogate this diversity, we micro-dissected developing human lungs along the proximal-distal axis during the late pseudoglandular stage and generated an integrated analysis of single-nucleus sequencing and spatial transcriptomics, creating a cellularly-resolved atlas of the lung. These rich datasets revealed positional niches and cellular heterogeneity along the proximal-distal axis, including the identification of a unique population of TP63^+^ basal cells, the primary stem cell of the airway, marked by expression of *LGR5* and *LGR6*. Analysis of the *LGR5+* basal cell niche and functional experiments with primary organoid models suggest a tonic level of WNT pathway activity in *LGR5^+^* basal cells that is potentiated by mesenchyme-derived R-SPONDIN. We found that basal cell self-renewal is enhanced by WNT activity, suggesting that the WNT pathway plays a previously unappreciated but critical role in airway stem cell maintenance during human development. These results enhance our fundamental understanding of the positional and cellular heterogeneity in the developing human lungs and begin to reveal unique niches that maintain homeostasis throughout the lung.

## INTRODUCTION

The lung epithelium plays a critical role in gas exchange, barrier protection, immunity, and electrolyte homeostasis. During development, morphogenesis and cell type differentiation occur in parallel^1,2^. The pseudoglandular stage occurs 5 to 17 weeks post-conception in humans and is defined by branching morphogenesis during which lung bud tips undergo successive bifurcations to create the airways. At the cellular level, the SOX9^+^ multipotent bud tip progenitors, which occupy the branching tips at the distal-most part of the airways, differentiate into basal cells and other airway cell types^3^. The KRT5^+^ TP63^+^ basal cells are a stem cell that are most abundant in the proximal cartilaginous airways, and play a role in airway homeostasis and repair^4,5^. Within the developing human lung, several single cell RNA-sequencing (scRNA-seq) datasets have revealed the diversity and heterogeneity of cell types within the epithelium^6–11^, including SCGB1A1/CC10^+^ secretory cells which produce secretoglobins, FOXJ1^+^ multiciliated cells involved in mucociliary clearance, MUC5AC^+^ goblet cells, FOXI1^+^ ionocytes, and CHGA^+^ pulmonary neuroendocrine cells (PNECs)^1^, and have led to the discovery of new epithelial types and subtypes, such as SCGB3A2^+^ SFTPB^+^ lower airway progenitor (LAP) cells^6,12^, and *GRP^+^* and *GRHL^+^* PNECs^7,8^. However, a spatially resolved atlas with cellular resolution in both the epithelium and mesenchymal compartments along the entire proximal-distal axis of developing human lung is lacking.

To characterize the complement of epithelial and mesenchymal cell types and niches with positional specificity, we performed micro-dissections to isolate four different regions; the trachea, large cartilaginous airways (bronchi), small non-cartilaginous airways, and distal/peripheral lung during the late pseudoglandular stage. We profiled gene expression and chromatin accessibility (snRNA-seq and snATAC-seq, 10X Multiomics), and used enriched genes to develop a custom probe-set for spatial transcriptomics (10X Xenium). By integrating snRNA-seq and high-resolution spatial information we observe both known, and previously unappreciated relationships between airway epithelial cell types, we define spatially-restricted mesenchymal niches and heterogeneity within the cartilage and heterogeneity along the proximal-distal axis within both epithelial and mesenchymal compartments. For example, within the proximal airways, we identified *LGR5^+^* basal cells and an epibasal KRT13^+^ population with cell bodies atop the basal cells. In the adult murine lung, Krt13^+^ cells have recently been described as a patch of cells in the trachea known as the ‘hillock’, and Krt13*^+^* cells are strongly induced following injury^13–15^. KRT13*^+^* cells are present but rare in the adult human trachea^14,15^; a similar cell type has not been described in human development.

Despite the well-investigated role of WNT signaling in stem cells of other organs^16^, within distal lung bud tips^3,17^ and in the alveolar compartment during development^18–20^, the WNT pathway has not been identified as an essential pathway for human basal cell function; human airway organoids can be expanded in media that lack activators of WNT signaling^6,12^. We observed *LGR5* and *LGR6* expression in airway basal cells, and expression of the LGR ligand *RSPO3* within the underlying mesenchymal cells of the airway niche. Given the ability of RSPO-LGR to potentiate WNT signaling^17,21–23^, we interrogated the requirement for RSPO-LGR-mediated WNT signaling in basal cell function. We established epithelium-enriched primary organoid cultures. The addition of exogenous R-SPONDIN protein to airway media enhanced expression of *LGR5* and *LGR6,* enhanced maintenance of the *LGR5^+^*basal cell, reduced cell differentiation, and improved colony formation efficiency. Conversely, addition of WNT pathway inhibitors reduced colony formation, suggesting that WNT is important for self-renewal. This is in contrast to the developing murine trachea where epithelial β-catenin deletion does not perturb basal cell development, and loss of Wnt signaling in the airway primarily affects the developing cartilage^24^. Together, this work adds essential spatial information for all cell types identified in the developing lung, defines a critical and human-specific role for WNT signaling in the developing airway, and has implications for *in vitro* modeling, directed differentiation, cellular therapy and transplant, and understanding and treatment of diseases of the airway.

## RESULTS

### Single-nucleus sequencing and spatial transcriptomics of micro-dissected developing human airways capture heterogeneity and regional specificity with cellular resolution

To investigate changes in cell composition along the proximal-distal axis of the developing human lung, we performed micro-dissections of fetal lungs with ages ranging from 101 to 132 days post conception (PCD). The lungs were dissected under the microscope using anatomic landmarks to separate distinct regions of trachea, primary bronchi, non-cartilaginous airways (NCA), and distal lung (Figure 1A, S1A). Immunofluorescence (IF) of TP63 (basal cells), SOX9 (cartilage, distal bud tips), and ECAD (epithelium) on representative sections was performed to confirm accuracy of the dissections, noting the presence of cartilage and basal cells along proximal airways, the absence of cartilage along the NCA, and the abundant bud tips distally (Figure S1B). The dissected tissue was used for several downstream analyses (Figure S1A), including single nuclear RNA-sequencing (n = 4; biological replicates from 101-120 PCD), paired single cell ATAC-sequencing by 10X Multiomics (n = 2; 102, 114 PCD samples), spatial transcriptomics by 10X Xenium (n = 2; 132, 137 PCD; 10X), histology, and the generation of primary region-specific epithelial organoid lines matched to specimens used for sequencing and additional for validation.

**Figure 1.**
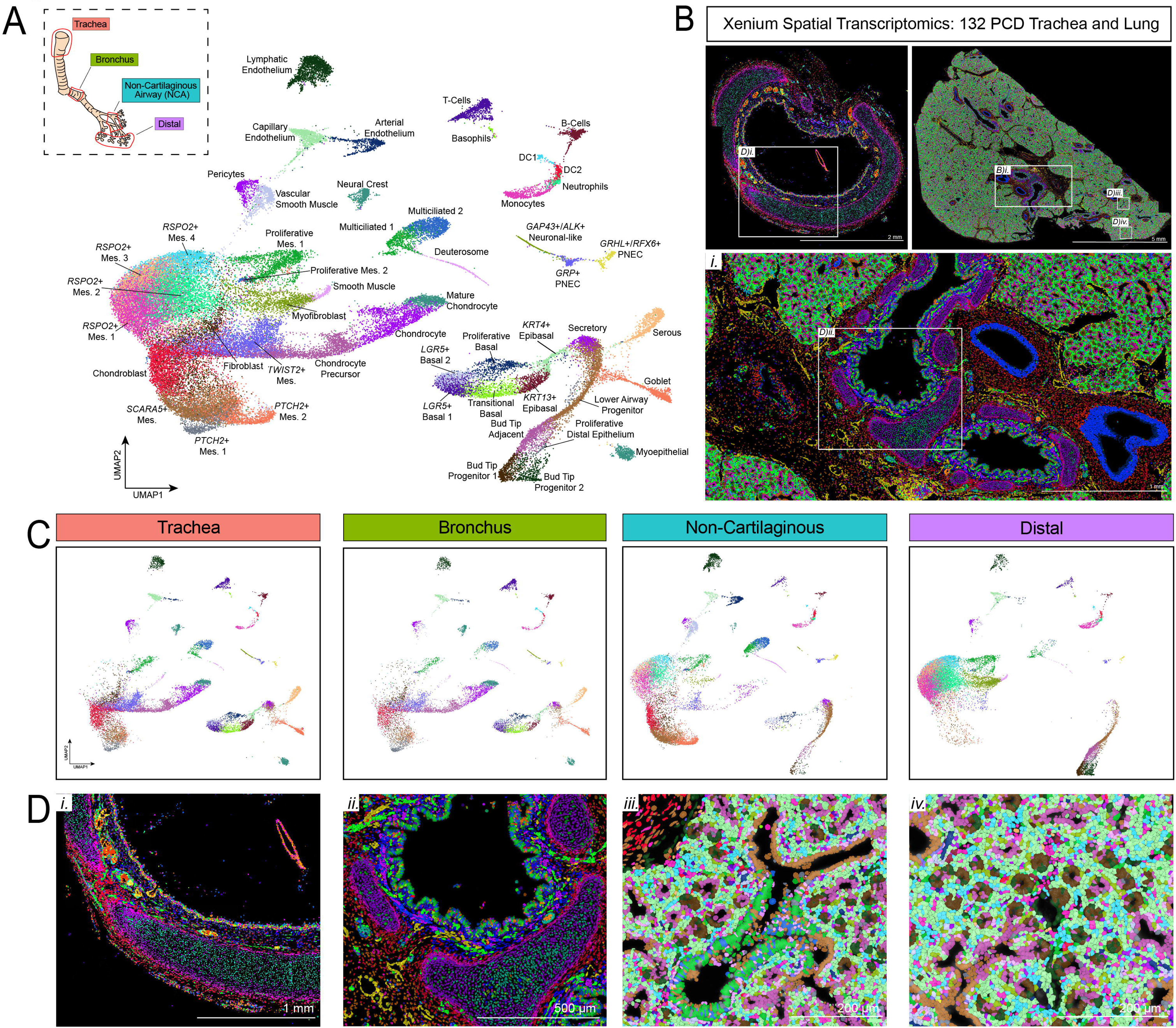
Generation of a positional atlas through sequencing and spatial transcriptomics of micro-dissected fetal lung regions. A. UMAP cluster plot of snRNA-sequencing data from human fetal lung micro-dissected regions including trachea, bronchus, non-cartilaginous airway, and distal regions, highlighted in inset. 4 biologically distinct samples were sequenced at PCD ages 101, 102, 115, 120. B. Xenium spatial transcriptomic image of 132 PCD human fetal trachea (Left) and lung (Right) with cells color coded by label transfer from identities defined in snRNA dataset (A). Insets for (B) and (D) highlighted accordingly. C. UMAP cluster plot from (A) split by dissected airway region. D. Xenium spatial transcriptomic images of 132 PCD human fetal trachea, bronchus, non-cartilaginous, and distal airway with cells color coded by label transfer from identities defined in snRNA dataset (A).

For snRNA-seq experiments, after ambient RNA correction and quality control (QC) filtering (Experimental Methods), 56,142 nuclei derived from all four regions and all four samples were included for analysis, which included Llouvain clustering and visualization by Uniform Manifold Approximation and Projection (UMAP) embedding (Figure 1A, Supplemental Table 1). We first assigned cells to 5 major cell classes, identifying epithelium (*CDH1*^+^), mesenchyme (*DCN*^+^), endothelium (*PECAM1*^+^), immune cells (*PTPRC*^+^), and neuronal cells (*SORCS1*^+^) (Figure S1C-D). Iterative sub-clustering and annotation was performed, resulting in 51 total cell types/states across all regions, including 19 epithelial, 19 mesenchymal, 7 immune, 3 endothelial, 2 neuronal, and 1 myoepithelial cluster (Figure 1A, S1E). We identified vessels, lymphatics and capillaries within the endothelial clusters, neural crest and neurons among the neuronal clusters, and lymphoid and myeloid lineages within the immune cells. Thus, our data appears to be representative of the diversity of cell types across tissue compartments of the lung (Figure 1A, S1E).

Based on genes enriched in each cell population defined in the snRNA-sequencing data (Figure S1E), we manually curated a customized 480-gene panel comprised of highly enriched markers (Supplemental Table 2) to perform 10X Xenium spatial transcriptomics on similarly aged samples. We analyzed the spatial transcriptomics data at both the level of cells and of individual genes. To assign cell identities in the Xenium data, we performed referenced-based mapping to the snRNA-seq data (Figure 1B). This allowed us to transfer the labels, assign identities to the cells in the spatial transcriptomic data, and visualize all cell types/states in tissue sections (Figure 1B, 1D). In general this approach identified cell types/states within the Xenium data that were concordant to cells present in the snRNA-seq data, providing confidence that this approach correctly assigned cell identities (Figure S2).

Given that micro-dissected regions of the lung were individually sequenced, this allowed us to computationally parse the snRNAseq into different regions, highlighting changes in cell composition along the proximal-distal axis (Figure 1C), and the spatial data confirmed the cell-type specification with added morphological and structural detail (Figure 1D). Broadly, these data confirmed proximal enrichment of basal cells and chondrocyte-lineage cells, distal enrichment of smooth muscle, lower airway progenitors, and bud tip progenitors in the distal lung with associated populations of mesenchyme that are highly specific to each region. For example, the proximal airway epithelium is supported by both a longitudinal band of smooth muscle and c-shaped cartilage rings, the lower airways are surrounded by smooth muscle and some cartilage, while the non-cartilaginous airways contain only smooth muscle (Figure 1D, S1F). One striking finding in our dataset is the presence of abundant and heterogeneous basal cell populations that are absent from published fetal lung atlases^7,8,11^, likely due to our deliberate inclusion of upper cartilaginous airway tissue.

### Heterogeneity of basal cell subtypes within the epithelium

To examine the epithelium with more granularity, we computationally extracted and re-clustered the epithelial compartment (Figure 2A-B), and found both expected and unexpected transcriptional heterogeneity within defined epithelial cell types, including 2 pulmonary neuroendocrine cells (PNEC) subtypes confirming recent reports by others^7,8^ (*GRP*^+^ PNEC, *GHRL*^+^PNEC, Figure S3A-D), 6 basal cell subtypes expressing canonical markers TP63 and KRT5 (*LGR5*^+^ Basal 1, *LGR5*^+^ Basal 2, Transitional Basal, *KRT13*^+^ Epibasal, *KRT4*^+^ Epibasal, Proliferative Basal), 2 bud tip progenitor subtypes (Bud Tip Progenitor 1, Bud Tip Progenitor 2), and 2 multiciliated cell clusters (Multiciliated 1, Multiciliated 2) (Figure 2A-B). Bud Tip Progenitor cluster 2 was distinguished by the onset of alveolar type 2 (AT2) markers (*C3, DMBT1, ABCA3*) and lower expression of bud tip progenitor markers (*TESC)*, likely reflecting an axis of nascent AT2 differentiation (Figure 2B).

**Figure 2.**
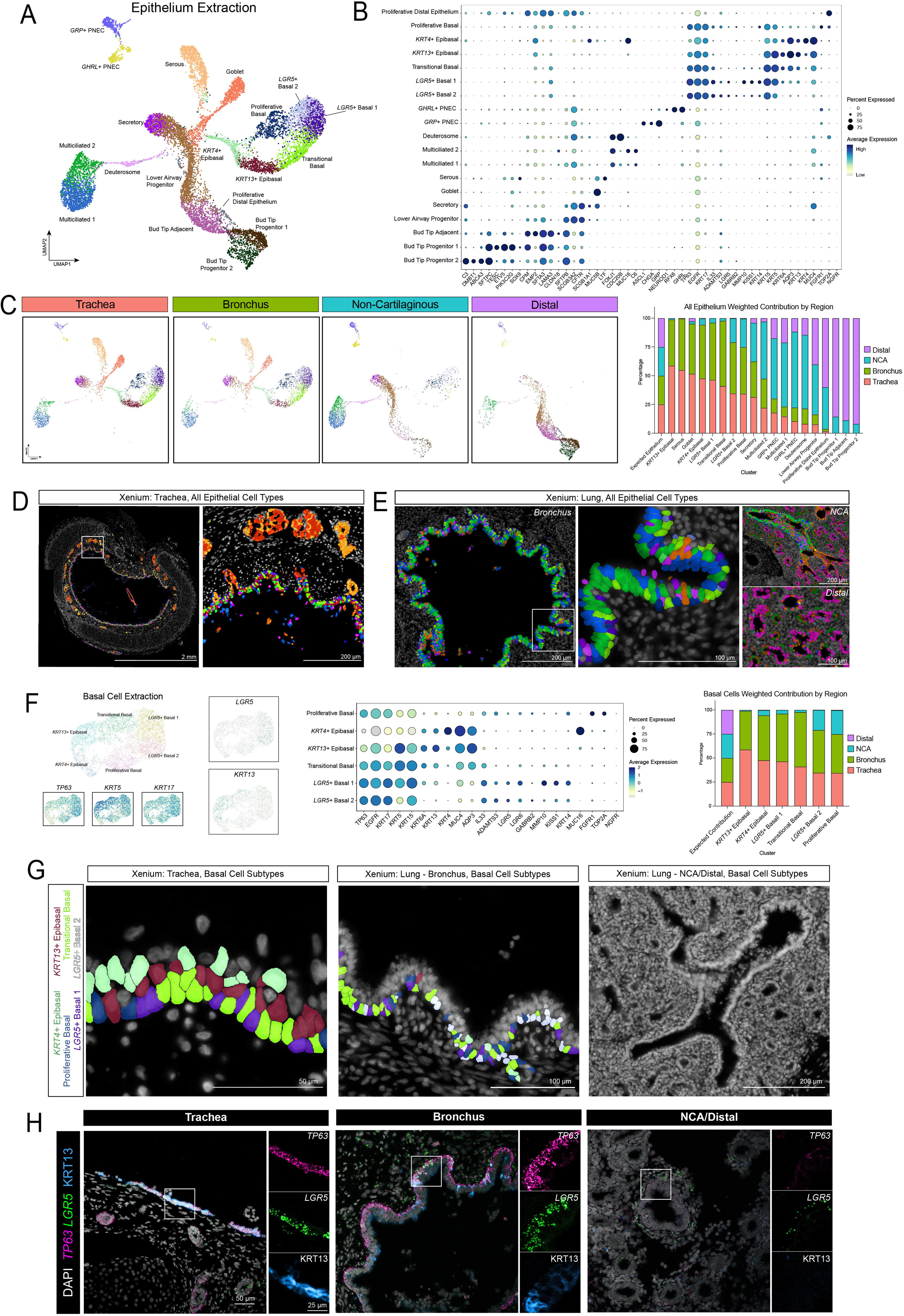
snRNA-seq and spatial analysis of micro-dissected lung regions uncovers regional epithelial heterogeneity and unique basal cell subtypes. A. UMAP cluster plot of snRNA-seq data from human fetal lung micro-dissected regions computationally extracted for epithelium only. B. Dot plot for expression of top genes defining each cluster of UMAP in (A). The dot size represents the percentage of cells expressing the gene in the corresponding cluster, and the dot color indicates log-normalized expression level of the gene. C. UMAP cluster plot from (A) split by dissected airway region. Distribution plot quantifies weighted percent of cells derived from each region. D. Xenium spatial transcriptomic image of 132d human fetal trachea with epithelial cells color coded by label transfer from identities defined in snRNA dataset (A). Inset highlighting the epithelium and submucosal gland cell types (colored), mesenchyme (grey). E. Xenium spatial transcriptomic image of 132d human fetal lung with epithelial cells color coded by label transfer from identities defined in snRNA dataset (A). Insets highlighting the epithelial cells (colored) in bronchus, NCA, and distal regions. F. UMAP cluster plot for computationally extracted basal cells, demonstrating 6 distinct clusters and basal cell sub-types (*LGR5+* Basal 1, *LGR5+* Basal 2, Proliferative Basal, Transitional Basal, *KRT13+* Epibasal, *KRT4+* Epibasal). Feature plots for canonical basal cell markers below (*TP63, KRT5, KRT17*). UMAP feature plots for sub-type defining genes (*LGR5*, *KRT13*). Dot plot of basal cell sub-type markers across all basal clusters. Distribution plot quantifies the weighted percent of each basal cell sub-type across 4 regions. G. Xenium spatial transcriptomic image of 132d human fetal trachea (left) and lung (middle, right)) with basal cells color coded by label transfer from identities defined in snRNA dataset (A). H. Representative fluorescence *in situ* hybridization (FISH) images with co-IF for *LGR5+* and *KRT13+* basal cell subtypes across trachea, bronchus, NCA and distal lung. Scale bar 50 μm, insets 25 μm.

Quantification with a weighted normalization to sample size contribution demonstrates the distribution of cell types and states along the proximal-distal axis (Figure 2C). 98.87% *KRT13*^+^ epibasal, 99.6% serous, and 95.15% goblet cells were derived from trachea and bronchus samples while 85.48% and 91.92% of bud tip progenitor clusters 1 and 2, respectively, were derived from distal samples, as expected. Secretory cells were found in a similar distribution across trachea (31.47%), bronchus (30.88%) and NCA (33.63%) samples. The majority of the Multiciliated 1 (55.73%) and Deuterosome clusters (64.15%) were isolated from the NCA. For Lower Airway Progenitor (LAP) cells, 83.59% were found in the NCA and distal samples, confirming our prior work describing LAP cells predominantly in the non-cartilaginous airways^12^. We also showed recently that two multiciliated states are present in the human lung^12^. Using barcoded lineage tracing in organoids, we demonstrated that one population arises from basal cells in the cartilaginous airways, distinguished by enriched *MUC16 expression,* and the other arises from LAP cells in the NCA region, distinguished by enriched *C6* expression^12^. In agreement with our prior findings, the current data showed *MUC16* expression enriched in the Multiciliated 2 population from which >50% of cells come from the cartilaginous airways, while the majority of Multiciliated 1 cells are enriched for C6 and are in the NCA and distal lung. The heterogeneity of epithelial cell distribution across the proximal-distal axis was supported by the Xenium data, where distinct cell types were visualized across trachea, bronchus, NCA and distal tissue regions (Figure 2D-E).

We were surprised by the heterogeneity of basal cell subtypes which had not been appreciated in other recent analyses. We computationally extracted and re-clustered the basal cells to better evaluate the predicted 6 subtypes (Figure 2F). Basal cell subtypes were identified with canonical markers *TP63, KRT5,* and *KRT17* which was expressed at similar levels across the subtypes (Figure 2F). Two of the subtypes, *LGR5^+^* Basal 1 and *LGR5^+^* Basal 2, are marked by expression of the genes *LGR5* and *LGR6* which encode the receptors for R-SPONDINs, agonists in the WNT pathway^16^. Within the *LGR5^+^* basal cells, *LGR5^+^* Basal 2 is enriched in the NCA (20.78% of *LGR5*^+^ Basal 2 cluster when evaluated in the weighted contribution) but is differentiated from the *LGR5*^+^ Basal 1 subset that is exclusive to the trachea and bronchi which also has higher expression of the genes *MMP10, KISS1,* and *MUC4* (Figure 2F). A third *TP63^+^ KRT5^+^ KRT17^+^* basal subtype is marked by *KRT13* expression, which is of note, as *Krt13* has been shown to mark a specific basal state within the mouse trachea^15^ and has only recently been found in adult human trachea^14,25^ (Figure 2F). To the best of our knowledge this is the first description of *KRT13^+^*basal cell state in development. A transitional basal cell exhibits intermediate expression between the *LGR5^+^* subtypes and the *KRT13^+^*subtype. A fifth subtype expresses *KRT4* (Figure 2F). Finally, the Proliferative basal cluster, which is *TOP2A*^+^, is found in a similar distribution across trachea (34.34%), bronchus (40.5%), and NCA (24.75%) samples (Figure 2F).

When we examined the spatial localization of the 6 subtypes in the trachea, we observed that the *LGR5^+^* basal cells were localized along the basal surface interspersed with Transitional or Proliferative basal cells. The *KRT13^+^* cells had nuclei and cell bodies just above the basal layer, which we refer to as the epibasal layer, while the *KRT4^+^* cells were localized more apically/lumenally within the pseudostratified epithelium of the developing trachea (Figure 2G, S3E). The organization of the subtypes within the developing epithelium suggests an axis of stemness that progresses from the basal layer (*LGR5^+^* cell) to the luminal side of the epithelium (Transitional-to-*KRT13^+^*-to-*KRT4^+^* basal cell) The complement of basal cells also changes moving proximally to distally. In the bronchus, there are fewer *KRT13^+^* and *KRT4^+^* epibasal cells. In the distal cartilaginous airways, there are proliferative basal cells and an enrichment of LGR5^+^ basal 2 cells. In the non-cartilaginous airways, basal cells are largely absent (Figure 2G, S3E). Fluorescence in situ hybridization (FISH) with co-immunofluorescence (co-IF) on human fetal lung sections was performed and validated the presence of these *LGR5*^+^ and KRT13^+^ basal subtypes in the trachea and bronchi, with limited presence in NCA and distal tissue, as well as the basal and epibasal locations within the pseudostratified proximal epithelium (Figure 2H). This was consistent with the spatial transcriptomic analysis when examined at the level of the individual probes (Figure S3F).

### Mesenchymal cell types are position- and niche-specific, expressing signaling receptors and ligands that form the basis for regional cell-cell communication networks

The mesenchymal clusters (Figure S1C-D) were computationally extracted and reclustered to evaluate their transcriptional signatures (Figure 3A-B). Many established mesenchymal populations were identified in this analysis as well as additional subclusters and specific niches along the proximal-distal axis. The recently described *RSPO2*-expressing mesenchyme was divided into 4 states, as we have previously suggested^17^, defined by varying levels of *RSPO2*, *PIEZO2*, and *FGFR4*. Our analysis also captured the transcriptional maturation of chondrocytes from an immature population we term ‘Chondroblast’ through a ‘Chondrocyte Precursor’ and to more terminal ‘Chondrocyte’ and ‘Mature Chondrocyte’ populations which have the highest expression of *COL11A1*, *CNMD*, *ACAN*, *COL9A1*, and *EPYC*. Other distinct mesenchymal cell types include cells defined by specific expression of *SCARA5*, *TWIST2*, or *PTCH2* (Figure 3A-B).

**Figure 3.**
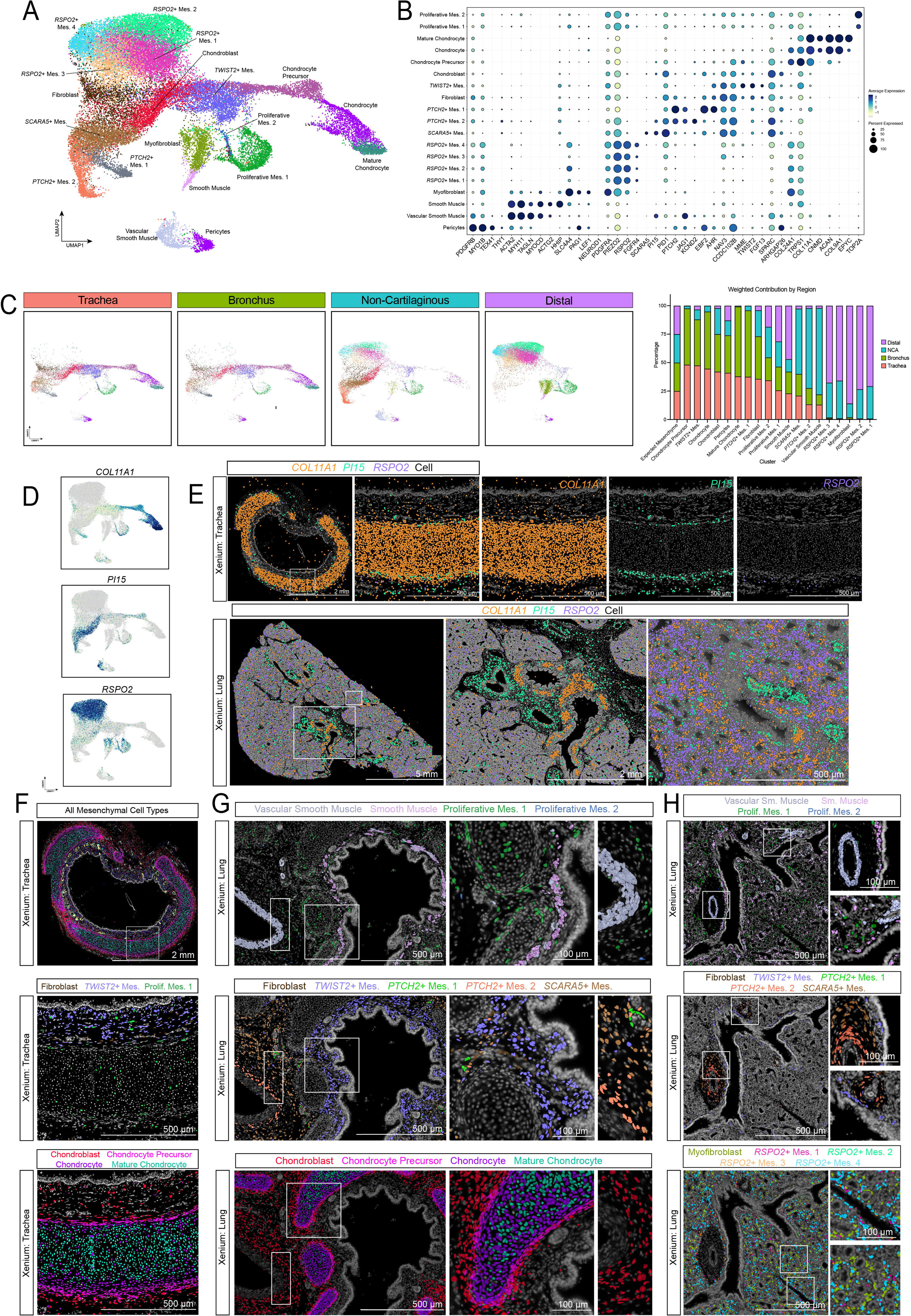
snRNA-seq and spatial analysis of micro-dissected lung regions uncovers mesenchymal heterogeneity and unique niches along the proximal-distal axis. A. UMAP cluster plot of snRNA-seq data from human fetal lung micro-dissected regions computationally extracted for mesenchyme only. B. Dot plot for expression of top genes defining each cluster of UMAP in (A). C. UMAP cluster plot from (A) split by dissected airway region. Distribution plot quantifies weighted percent of cells derived from each region. D. UMAP feature plots for genes encompassing proximal-to-distal regionality: *COL11A1* in proximal (trachea/bronchus); *PI15* in NCA; *RSPO2* in distal. E. Xenium spatial transcriptomics image of *COL11A1*, *PI15*, and *RSPO2* transcript expression in 132d human fetal trachea (top) and lung (bottom). F. Xenium spatial transcriptomics image of 132d human fetal trachea for mesenchymal subtypes color coded by label transfer from identities defined in snRNA dataset (A). Insets highlight subsets of mesenchymal cell types including Fibroblast, *TWIST2*+ Mesenchyme, Proliferative Mesenchyme 1, and the chondrocyte lineages. G. Xenium spatial transcriptomics image of 132d human fetal bronchus for specific mesenchymal subtypes color coded by label transfer from identities defined in snRNA dataset (A). Insets share a scale bar 100 μm. H. Xenium spatial transcriptomics image of 132d human fetal NCA and distal airways for specific mesenchymal subtypes color coded by label transfer from identities defined in snRNA dataset (A). Insets share a scale bar 100 μm.

When the populations are evaluated by their regional specificity (Figure 3C), the chondrocyte lineage is predominantly derived from the trachea and bronchi (90.1%), and the distal *RSPO2* populations are isolated from the NCA and distal samples (98.8%). The *TWIST2*^+^ population is proximal-specific with 87.67% derived from trachea and bronchus, while the majority of the *SCARA5*^+^ population is from NCA (58.62%). The two *PTCH2*^+^ populations are also defined by their regional specificity, with *PTCH2^+^* Mesenchyme 1 derived from proximal samples and *PTCH2*^+^ Mesenchyme 2 from NCA. We also noticed that three markers could be used to coarsely define groups of mesenchymal cell types along the proximal-distal axis. *COL11A1*, predominantly found in cartilage, is highly expressed in the proximal regions of mesenchymal tissue, *PI15* is expressed throughout the proximal and intermediate regions, and *RSPO2* broadly captures all distal mesenchyme with the exception of pericytes (Figure 3D-E). When examined more closely, there are small populations of *PI15* along the cartilaginous rings in trachea tissue with more broad expression throughout the blood vessels (Figure 3E, S4A).

We also leveraged our spatial data to examine the localization of mesenchymal populations defined by our snRNA-sequencing (Figure 3F), once again using reference-mapping to make cell identifications. In the trachea sample we observed Proliferative Mesenchyme populations, Fibroblasts and *TWIST2*^+^ Mesenchyme localized between epithelium and cartilage. Interestingly within the *TWIST2*^+^ Mesenchyme population, there were some rounded cells near the epithelium and elongated cells closer to the cartilage visualized in our spatial transcriptomic data suggesting additional heterogeneity (Figure 3F). Cells we assigned to the chondrocyte lineage (Chondroblast, Chondrocyte Precursor, Chondrocyte, Mature Chondrocyte) were organized in a radial fashion, with Chondroblasts occupying the perimeter of cartilage rings, followed by inward layering of Precursors, Chondrocytes and Mature chondrocytes in the center (Figure 3F, S4B). *PTCH2^+^* Mesenchyme 1 was localized around submucosal glands in the trachea and bronchi. *PTCH2^+^* Mesenchyme 2, was enriched in NCA samples where it localized directly adjacent to vascular smooth muscle at larger vessels (Figure 3G.H). *SCARA5^+^* mesenchyme was localized to the interstitial regions (Figure 3G). Distally, we were able to appreciate the intimate association among the bud tip epithelium, *RSPO2^+^* Mesenchyme, the Myofibroblasts, developing Capillaries, and Lymphatics (Figure 3H, S4C-D).

### The proximal airway epithelium is predicted to participate in autocrine WNT signaling that is modulated by adjacent mesenchyme

Given the surprising basal cell heterogeneity, the abundance of a previously unappreciated *LGR5^+^* basal cell occupying the most ‘basal’ position within the pseudostratified epithelium, and the known role of LGRs for potentiating WNT signaling, we sought to understand the signaling interactions between the proximal epithelium and supporting mesenchyme. We used a coarse annotation of the trachea and bronchus epithelial and mesenchymal cells (Figure 4A-B) to focus on broad signaling patterns across the two compartments and interrogated cell-cell communication using the CellChat^26^ package (Figure S5A). CellChat predicted that the *LGR5*^+^ basal cell can signal to itself in an autocrine fashion in addition to other epithelial clusters such as the epibasal and proliferative basal cells; CellChat also predicted the mesenchyme as also a strong source of WNT signal to the epithelial cells. Because the genes encoding R-SPONDINs (*RSPO*) are not included in CellChat’s database of signaling interactions, we also individually examined the expression of WNT signaling pathway components (Figure 4C-D, S5B). *FZD* receptors 1, 2, and 4 are broadly expressed in mesenchymal cell types while *FZD3 and FZD6* are more specific to the epithelium. The R-SPONDIN receptor genes *LGR5* and *LGR6,* which potentiate WNT signaling, are restricted to basal cell populations. Regarding ligands, *WNT3A, WNT4*, and *WNT7B* are specifically expressed by the basal cell populations, *WNT2B* is expressed by both the epithelium and mesenchyme, while *RSPO1-3* is only in the mesenchyme. When we examine the expression of RSPOs comparing proximal to distal lung, we find that while *RSPO1-3* are all expressed to an extent in the mesenchyme of the trachea and bronchus, *RSPO3* is the most highly enriched in the proximal airway (Figure 4C-D, S5B). Based on these analyses of expression patterns, we therefore hypothesized that basal cells are able to sustain a certain degree of active WNT pathway activity through autocrine WNT-FZD signaling but that the mesenchyme sends additional potentiating RSPO signals to LGR5 and LGR6.

**Figure 4.**
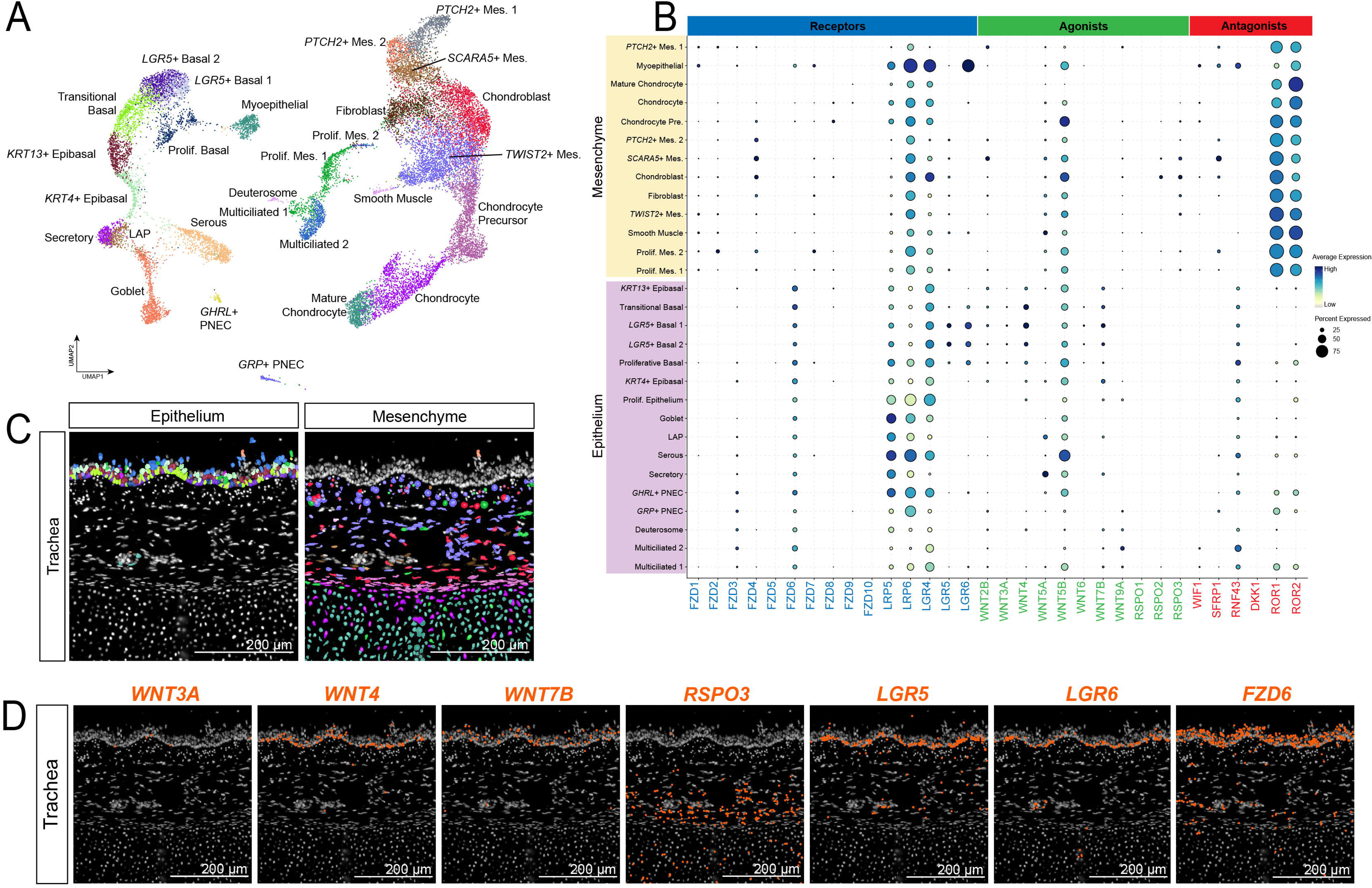
The proximal airway epithelium is predicted to participate in autocrine WNT signaling via WNT-FZD receptor-ligand interactions and is modulated by adjacent mesenchyme which express the R-SPONDINs. A. UMAP cluster plot of snRNA-seq data from human fetal lung, trachea and bronchus samples only, for computationally extracted epithelium and mesenchyme clusters. B. Dot plot for expression of WNT receptors, agonists and antagonists across mesenchymal and epithelial clusters annotated in (A). C. Xenium spatial transcriptomics image of 132d human fetal trachea for epithelial (left) and mesenchymal (right) subtypes identified in trachea, color coded by label transfer from identities defined in snRNA dataset (A). Region reference for images in (D). D. Xenium spatial transcriptomics image of WNT signaling components (*WNT3A*, *WNT4*, *WNT7B*, *RSPO3*, *LGR5*, *LGR6*, *FZD5*) transcript expression in 132d human fetal trachea.

### Primary epithelial organoids maintain regional heterogeneity

To functionally interrogate the hypothesis that RSPO-LGR interactions potentiate WNT signaling in basal cells, and to investigate the epithelial heterogeneity identified in sequencing and spatial transcriptomics data, we developed primary organoid models from the four regions of interest (Figure 1A, 5A). Details of organoid generation, validation, and culture conditions can be found in ‘Experimental Methods’ and were described previously by us^12^. Briefly, minced tissue from trachea, bronchus, NCA and distal lung tissue were embedded in Matrigel and established in our previously published airway media^6^ for 1 week before being dissociated to single cells, re-embedded in Matrigel for continued culture, and allowing for epithelial organoid formation. Organoids from all four regions were analyzed by snRNA-sequencing after passage 2 of organoid generation. After ambient RNA removal and QC filtering, 18,185 cells were analyzed and visualized with a UMAP embedding (Figure 5B, S6A). Clustering identified 13 clusters (Figure 5B, S6A), and region-by-region analysis confirmed that the regional specificity of these clusters is captured in culture (Figure S6B-C). Clusters 0, 2, and 3 were predominantly isolated from trachea, while clusters 4, 5, 6, 8, and 10 were notably derived from NCA and distal tissue (Figure S6B-C).

**Figure 5.**
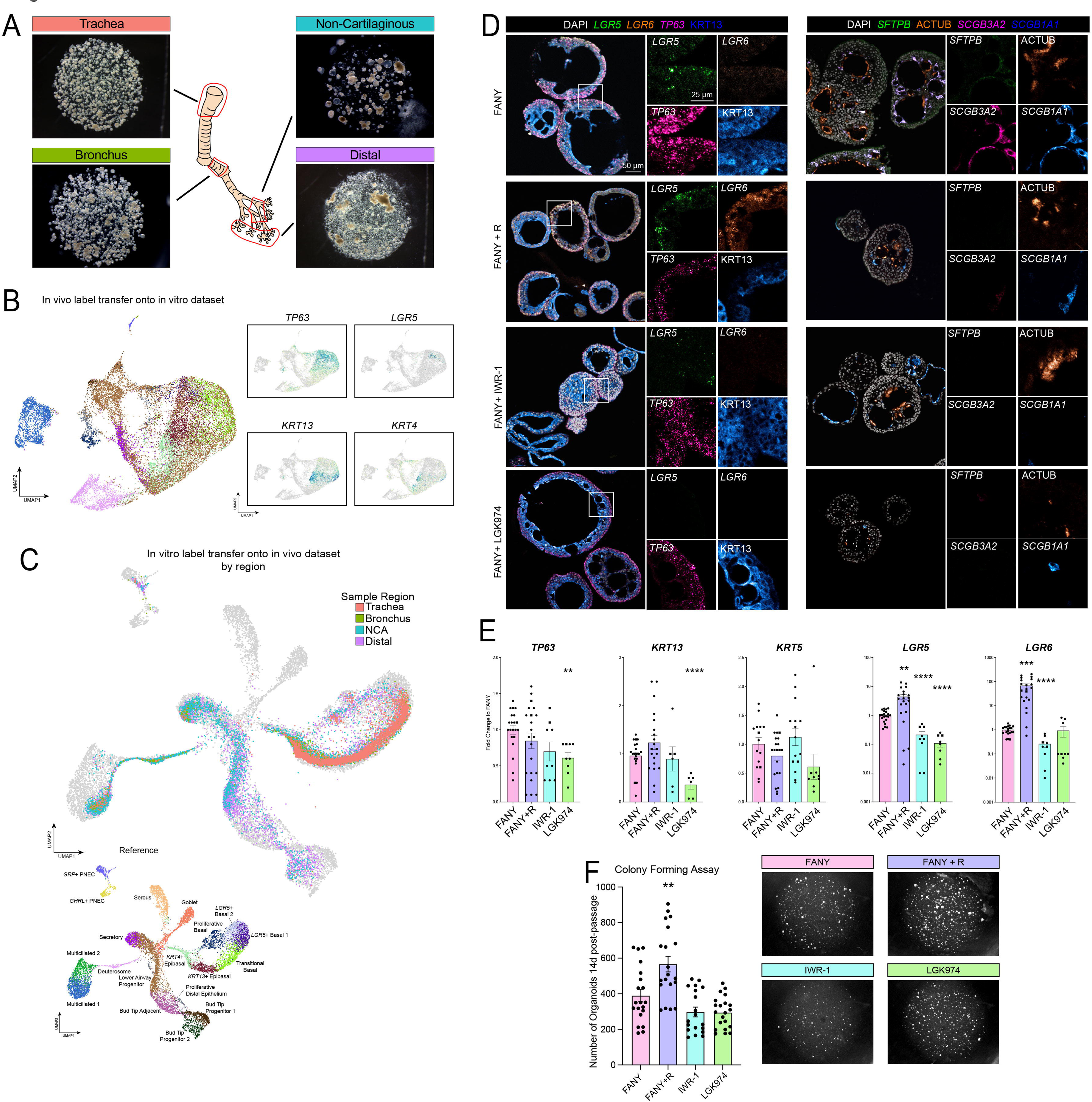
Potentiation of tonic WNT signaling with the addition of R-SPONDIN supports the maintenance of *LGR5+* basal cells in culture. A. Representative brightfield images of organoid cultures derived from 4 regions of dissected primary tissue. B. UMAP cluster plot of snRNA-seq data from organoid lines (n=1) combining all 4 regions. Colored by label transfer of *in vivo* snRNA-seq epithelial dataset (Figure 2A). Feature plots for basal cell markers (*TP63*, *LGR5*, *KRT13*, *KRT4*). C. UMAP cluster plot of snRNA-seq data from organoids (colored) label transferred onto *in vivo* epithelial dataset (grey). Inset of reference UMAP from Figure 2A. D. Representative FISH with co-IF of organoids before (FANY) and after (FANY+R) stimulation of WNT with RSPO1 in culture. Additional conditions of WNT inhibitors (IWR, LGK). Markers include basal cell-types (*LGR5, LGR6, TP63,* KRT13) and other airway cells (*SFTPB, SCGB3A2, SCGB1A1,* AcTUB). E. RT-qPCR of basal cell markers in WNT modulation experiment. This quantification was performed on a minimum of three separate biological replicates with three technical replicates each. Error bars represent standard error of the mean. Statistical tests were performed by Welch ANOVA test with Dunnett’s T3 multiple comparisons. P-values ** < 0.005, *** < 0.0005, **** < 0.0001. F. Quantification of colony formation 14 days after dissociation to single cells from organoid lines (n = 2). This quantification was performed on four separate experimental replicates with a minimum of three technical replicates each. Error bars represent standard error of the mean. Statistical tests were performed by Welch ANOVA test with Dunnett’s T3 multiple comparisons. P-value ** = 0.01.

To annotate the cell types captured among the organoid cultures, we performed reference-based mapping from our *in vitro* cells to our *in vivo* snRNA-sequencing epithelial subset (Figure 5B-C, S6D-E). This approach provided an unbiased method to classify the cell types present *in vitro* that takes into account common variable features, otherwise known as anchors, for each dataset. Performing this analysis on integrated data from all organoids and then examining cell identity assignments depending on which region the organoid was derived from revealed that organoids maintained cellular compositions consistent with our *in vivo f*indings (Figure 5C). To better highlight which populations were or were not captured in the *in vitro* organoids, we plotted the *in vitro* cells onto the reference *in vivo* UMAP. This highlighted that organoids from the trachea and bronchi were highly enriched for *KRT13^+^*basal cells while organoids from the NCA were highly enriched for LAP cells, corresponding to their proximal-distal region origin (Figure 5B-C). It was also apparent that the cultures strikingly lack the abundant *LGR5*^+^ basal subtypes despite retaining other basal cell states (Figure 5B-C).

### Potentiation of tonic WNT signaling with the addition of R-SPONDIN supports the maintenance of *LGR5^+^* basal cells

Our standard airway media consists of a base media and supplements along with <u>F</u>GF10, dual SMAD inhibitors <u>A</u>8301 and <u>N</u>oggin, and the ROCK inhibitor <u>Y</u>-27632 (<u>FANY</u>)^6^ (Experimental Methods). Given that our basal media does not contain an exogenous WNT ligand or agonist and our cultures are epithelial without a mesenchymal component, it is likely that our culture conditions do not support the level of WNT activity required to maintain the *LGR5^+^* basal cell population *in vitro,* as is suggested by snRNA-seq label transfer of airway organoids (Figure 5C). To recapitulate the mesenchyme-derived RSPO signaling axis, we added <u>R</u>SPO1 which can act as a ligand for both LGR5 and LGR6, into our standard FANY media (<u>FANY</u> + <u>R</u>).

Primary organoids derived from the trachea were grown in FANY or FANY + R for 3 weeks and analyzed by FISH/IF and RT-qPCR (Figure 5D-E, S6F). We found that the addition of RSPO1 led to a significant *LGR5* and *LGR6* induction, while *TP63* and *KRT5* levels were similar between FANY and FANY + R (Figure 5D-E). By FISH/IF, we observed that while in FANY media there are few *LGR5^+^* or *LGR6^+^* cells, FANY + R media rescues the *LGR5* and *LGR6* expression within *TP63^+^* cells on the basal side of the organoid epithelium (Figure 5D). Conversely, the addition of the WNT inhibitors IWR-1, which leads to β-Catenin degradation, or LGK974, which inhibits porcupine and prevents all endogenous WNT secretion, led to a decrease in *LGR5* and *LGR6* expression beyond the low level in our standard conditions (Figure 5D-E). When we examined markers of airway differentiation by IF, we saw that RSPO1 addition led to decreased expression of markers of differentiated cell types like *SCGB1A1* (secretory), *FOXJ1* (ciliated), and *MUC5AC* (goblet) (Figure 5D, S6F). These data suggest that RSPO-LGR signaling potentiates WNT activity, maintains *LGR5* and *LGR6* expression, and blocks differentiation, while lower WNT activity leads to even lower *LGR5* and *LGR6* expression and enhanced differentiation. To further interrogate how the addition of RSPO1, IWR-1 or LGK974 influences basal cell function, we performed an organoid colony forming assay on n = 2 different biological specimens. For this experiment, tracheal organoids were grown in FANY in early passages, and starting from passage 3/4, organoids were dissociated into single cells every 14 days, counted, and 3,000 cells were plated into a droplet of Matrigel, switched to media containing RSPO1, IWR-1 or LGK974. They were allowed to recover for 14 days, and the number of organoids formed were quantified. We found that RSPO1 increases the efficiency of organoid colony formation (Figure 5F) while WNT inhibitor treatments lead to a trend in decreased colony formation efficiency, suggesting that WNT is functionally important for basal cell potency (Figure 5F).

## DISCUSSION

Here, we build on several prior studies by our group and others^6–8,11,25^ who have interrogated human lung development at single cell resolution, mapping out a blueprint consisting of the full complement of heterogeneity at cellular resolution across the proximal-distal axis of the developing human lung. We attribute the development of this powerful resource to our unique experimental approach and multimodal analyses. We deliberately focused our tissue collection on different regions of the airway, micro-dissecting the fetal lung from the proximal trachea, across anatomically diverse (cartilaginous vs. non-cartilaginous) regions of the airways to the distal lung buds. Integrating single-nuclear RNA sequencing with our custom spatial transcriptomic analysis has been instrumental for assessing the cellular and signaling heterogeneity and has made possible the identification of *LGR5*^+^ basal cells, the *RSPO3*^+^ niche cells, and for dissecting cell-cell interactions. Our data will be an important community resource for precious human tissues, providing access to a broad base of scientists that can use these data for hypothesis-generation without the need for additional computational manipulation.

The WNT pathway plays an essential and conserved role in development and stem cell homeostasis across multiple organs^16^. While we have examined the role of WNT activity in the distal bud tip during human lung development^17^ and Wnt signaling is important for secretory and ciliated cells in the murine airway^27^, for primary human adult cells in culture^28–31^, and the airway mesenchyme in mouse and pig^32–34^, we believe that this is the first report showing a requirement for WNT in the developing airway epithelium. In contrast to the developing human lung, within the mouse lung during development, genetic experiments that block secretion of WNT ligands from the epithelium show a primary defect in the cartilage^24^, and genetic deletion of β-catenin from the entire epithelium does not disrupt basal cells^24^, suggesting that WNT/β-catenin signaling is dispensable for basal cell maintenance or regulation. We find that the *LGR5^+^* basal cell is abundant in human lung development, lining the fetal proximal airways, the cell bodies positioned most basally within the pseudostratified epithelium. Our functional organoid experiments suggest that WNT pathway activity is required for self-renewal and maintenance of the *LGR5^+^* basal cell and that while the epithelium maintains a tonic level of activity through autocrine WNT-FZD signaling, potentiation occurs from RSPO secretion from the adjacent mesenchyme. WNT signaling can be classified into a canonical, β-catenin-dependent or a non-canonical, β-catenin-independent pathway^16^. Our data with WNT inhibitors supports a role for the canonical pathway in fetal basal cells.

In addition to the *LGR5*^+^ basal cell, we identify a similarly abundant *KRT13*^+^ epibasal cell. These cells have been previously described in the adult mouse airway within a structure deemed the “hillock” which consists of islands of stratified squamous epithelium existing amidst the pseudostratified epithelium^15^. The hillock Krt13+ basal cells are speculated to be involved in wound-healing, where they replace denuded epithelium, form an injury-resistant shield and are eventually able to differentiate into the major cell types of the airway^14^. Hillock structures also seem to be present in adult human trachea but, unlike in mouse, low-expressing KRT13^+^ suprabasal cells also seem to be interspersed within the pseudostratified epithelium^14^. In the developing airway we have yet to appreciate a squamous structure reminiscent of the hillock, but instead find the *KRT13*^+^ basal cells are interspersed among the *LGR5^+^* basal cells. KRT13^+^ cells are plentiful in our positional airway organoids derived from the proximal airways even in the absence of WNT agonist. Whether they represent a transit-amplifying population or a similarly multipotent progenitor in comparison to the *LGR5^+^* basal cells remains unclear. Moreover, whether these KRT13^+^ cells eventually condense into hillock-type structures, and when this might occur later in development is unknown. If in adulthood the hillock exists to form an injury-resistant barrier and as a reserve source of airway cells, it is unclear the function of the structure is required during development but it may perhaps play a role in rapidly colonizing the growing trachea with epithelial cells.

Cellular therapy using directed differentiation of pluripotent stem cells into airway progenitors is an active area of interest for the treatment of airway diseases such as Cystic Fibrosis^35^. Our data along with other adult atlases will be an important reference with which to benchmark airway cells. Moreover, our work with primary fetal airway organoids suggests that the addition of WNT ligands and agonists will be important for *in vitro* expansion and proper regional patterning. Whether the addition of WNT ligands and agonists *in vitro* or *in vivo* improves the efficacy of cellular transplant to the airways is an area that we are actively investigating and may have implications for the efficacy of stem cell therapy. Finally, the *LGR5^+^* basal cell is rare in the homeostatic adult airway^36^. As Wnt signaling is also absent from the murine airway at homeostasis, but is highly induced following injury^27,29,37–40^, it is also interesting to speculate that the low level of WNT activity in the adult human airway reflects slow turnover during homeostasis, but that injury and the need for tissue repair and regeneration may require WNT signaling, mediated in part by RSPO-LGR interactions. Thus, whether WNT pathway activity is important for repair after airway injury or plays a role in human lung disease remains to be explored.

## MATERIALS AND EXPERIMENTAL METHODS

### Human Lung Tissue

Human lung tissue research was reviewed and approved by The University of Michigan Institutional Review Board (IRB). Normal, de-identified human lung tissue was obtained from the University of Washington Laboratory of Developmental Biology. Tissue was shipped overnight in UW-Belzer’s solution (Thermo Fisher, Cat#NC0952695) on ice and was processed for experiments or fixation within 24h.

### Positional Airway Organoid Establishment and Culture

Organoid establishment was carried out as previously described^12^. Proximal trachea, bronchus, non-cartilaginous airway, and distal tissue was dissected from tissue using a scalpel and forceps and subsequently minced for 2 min into small chunks (<1mm^3^). Tissue was washed with 1X PBS, spun down at 300g for 3 min at 4°C, resuspended in Matrigel, replated in 20 µL droplets and allowed to solidify. Cultures were fed with 0.5 mL FANY media (ref) for 1 week, changing media every 3-4 days, after which they were passaged using TrypLE (Invitrogen, Cat#12605010), for 30 min with mechanical dissociation every 10 min. Dissociation was quenched using DMEM/F-12 (Corning, Cat#10-092-CV) after which cells were spun down at 300g for 3 min at 4°C, resuspended in Matrigel and fed with 0.5 mL FANY media. Cultures were passaged by single cell dissociation using TrypLE every 2 weeks.

### Organoid Forming Efficiency Assay

Airway organoids were collected using a cut P1000 pipette tip and transferred to a 15 mL conical tube. Organoids were centrifuged at 500g for 5 min at 4°C. After centrifugation, the media was aspirated, and ∼5 mL of cold 1X PBS was added. The organoids were vigorously pipetted to remove as much Matrigel as possible, then centrifuged again at 500g for 5 min at 4°C. Excess liquid and remaining Matrigel were removed. Organoids were digested into single cells using 3-4 mL of TrypLE (Invitrogen, Cat#12605010), depending on organoid size, and incubated at 37°C for 60-90 min. During this incubation, mechanical dissociation was performed every 5-10 minutes by pipetting up and down with a P1000 tip and/or P20 tip on top of a P1000 tip until a single-cell suspension was achieved. Trypsinization was quenched with warm DMEM/F-12 (Corning, Cat#10-092-CV). Using 1 mL DMEM/F-12 for prewetting and 2 mL for rinsing, cells were filtered through a 40 μm Cell Strainer (ThermoFisher, Cat#08-771-1) and centrifuged at 500g for 5 min at 4°C, followed by aspiration of the supernatant. Depending on pellet size, cells were additionally washed with 1-2 mL of cold 1 PBS, centrifuged at 500g for 5 min at 4°C and supernatant was removed. The cell pellet was resuspended in 100-500 μL of 1X PBS and counted using a Countess™ Automated Cell Counter (Invitrogen, Cat#C10227) with 0.4% Trypan Blue Stain (Invitrogen, Cat#T10282). After counting, the desired number of cells was transferred to a 1.5 mL mini-centrifuge tube and centrifuged at 500g for 5 min at 4°C. The supernatant was aspirated. Cells were resuspended in Matrigel (Corning, Cat#354234) at a concentration of 75 cells/μL. 5x 40 μL droplets of Matrigel were plated into each well of a 6-well tissue culture plate (ThermoFisher, Cat#FB012927). After 3 min at RT, the plate was inverted and placed in a 37°C incubator for 15 minutes. Organoids were fed every 3-4 days with the following media conditions: FANY+DMSO (1000X, Sigma-Aldrich, Cat#D2650), FANY+R+DMSO (1000X), FANY+IWR1 (5 μM, Fisher Cat#50-101-4191), and FANY+LGK974 (250 nM, MedChemExpress, Cat#HY-17545). After two weeks, organoids were imaged using an Olympus SZX16 microscope, and their number and size were quantified using the Python scikit-image package with the watershed algorithm.

### Tissue Processing and Staining

All sectioned fluorescent images were taken using a Nikon AXR confocal microscope. Acquisition parameters were kept consistent for images in the same experiment and all post-image processing was performed equally on all images in the same experiment. Images were assembled in Adobe Photoshop CC 2024.

#### Tissue Processing

Tissue was immediately fixed in 10% Neutral Buffered Formalin (NBF) for 24h at room temperature on a rocker. Tissue was then washed 3x in UltraPure DNase/RNase-Free Distilled Water (Thermo Fisher, Cat#10977015) for 15 min each and then dehydrated in an alcohol series of concentrations dehydrated in UltraPure DNase/RNase-Free Distilled Water for 1h per solution: 25% Methanol, 50% Methanol, 75% Methanol, 100% Methanol, 100% Ethanol, 70% Ethanol. Dehydrated tissue was then processed into paraffin blocks in an automated tissue processor (Leica ASP300) with 1 hr solution changes. 5 (FISH) or 7 (IF) µm-thick sections were cut from paraffin blocks onto charged glass slides. For FISH, microtome and slides were sprayed with RNase Away (Thermo Fisher, Cat#700511) prior to sectioning (within one week of performing FISH). Slides were baked for 1 hr in 60°C dry oven (within 24 hr of performing FISH). Slides were stored at room temperature in a slide box containing a silica desiccator packet and the seams sealed with paraffin.

#### Immunofluorescence (IF) Protein Staining on Paraffin Sections

Tissue slides were rehydrated in Histo-Clear II (National Diagnostics, Cat#HS-202) 2x for 5 min each, followed by serial rinses through the following solutions 2x for 3 min each: 100% EtOH, 95% EtOH, 70% EtOH, 30%EtOH, and finally in double-distilled water (ddH2O) 2x for 5 min each. Antigen retrieval was performed using 1X Sodium Citrate Buffer (100mM trisodium citrate (Sigma, Cat#S1804), 0.5% Tween 20 (Thermo Fisher, Cat#BP337), pH 6.0), steaming the slides for 20 min, followed by cooling and washing quickly 2x in ddH2O and 2x in 1X PBS. Slides were incubated in a humidified chamber at RT for 1 hr with blocking solution (5% normal donkey serum (Sigma, Cat#D9663) in PBS with 0.1% Tween 20). Slides were then incubated in primary antibody diluted in blocking solution at 4°C overnight in a humidified chamber. Next, slides were washed 3x in 1X PBS for 5 min each and incubated with secondary antibody with DAPI (1µg/mL) diluted in blocking solution for 1h at RT in a humidified chamber. Slides were then washed 3x in 1X PBS for 5 min each and mounted with ProLong Gold (Thermo Fisher, Cat#P36930) and imaged within 2 weeks. Stained slides were stored in the dark at 4°C. All primary antibody concentrations are listed in Supplementary Table 4. Secondary antibodies, raised in donkey, were purchased from Jackson Immuno and used at a dilution of 1:500.

#### Fluorescence in situ hybridization (FISH)

The FISH protocol was performed according to the manufacturer’s instructions (ACDbio, RNAscope multiplex fluorescent manual) with a 5-minute protease treatment and 15-minute antigen retrieval. For IF co-staining with antibodies, the last step of the FISH protocol was skipped and instead the slides were washed 1x in 1X PBS followed by the IF protocol above. A list of probes and reagents can be found in Supplementary Table 4.

IF and FISH stains were repeated on at least 3 independent biological replicates/experiments and representative images are shown.

#### Tissue Preparation for Xenium Spatial Transcriptomics Analysis

Excess paraffin was trimmed from the paraffin blocks, which were subsequently submerged in an ice bath for at least 30 minutes prior to sectioning. To prepare for section collection, the slide warmer was preheated to 42°C and used to pre-warm positive-charged microscope slides topped with molecular biology grade water (Corning, Cat# 46-000-CM). The blocks were then sectioned at a thickness of 5μm, and serial sections were collected on the prepared slides. After allowing sections to flatten out, the water was removed, and sections were examined under the 4X and 10X objectives of a biological microscope to identify the optimal regions of the samples. Using a single edge razor blade, the corresponding area of the paraffin block was lightly scored to obtain tissue sections small enough to fit into the 10.45mm by 22.45mm sample placement area of the Xenium slide. Xenium slides were removed from -20°C storage and allowed to come to room temperature 30 minutes prior to undergoing the same preparation as the microscope slides. Once the selected sections were placed on the Xenium slide, they were examined under the biological microscope again to ensure that the fiducial frame was not obscured by any paraffin or tissue. Following water removal, sections were allowed to dry on the slide warmer for 3 hours before slides were moved to a microscope slide jar for transportation to the Advanced Genomics Core.

### RNA extraction, cDNA, RT-qPCR

Each analysis includes at least three technical replicates from three separate biologic tissue lines. mRNA was isolated using the MagMAX-96 Total RNA Isolation Kit (Thermo Fisher, Cat#AM1830). RNA quality and yield was measured on a Nanodrop 2000 spectrophotometer just prior to cDNA synthesis. cDNA synthesis was performed using 100ng RNA per sample with the SuperScript VILO cDNA Kit (Thermo Fisher, Cat#11754250). RT-qPCR was performed on a Step One Plus Real-Time PCR System (Thermo Fisher, Cat#42765592R) using QuantiTect SYBR Green PCR Kit (Qiagen, Cat#204145). Primer sequences can be found in Supplementary Table 4. Gene expression as a measure of arbitrary units was calculated relative to Housekeeping gene (18S) using the following equation:

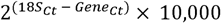

### Quantification, Statistical Analysis

For RT-qPCR analysis, biologic replicates containing 3 wells of organoids, each containing 10-60 organoids (technical replicates) were collected. All statistical analyses were performed in GraphPad Prism Software. See Figure Legends for the number of replicates used, statistical test performed, and the p-values used to determine the significance for each separate analysis.

### snRNA-seq/ATAC-seq experiments

#### Single-nucleus isolation and permeabilization

Preparation was performed as described previously^41^ in accordance with 10X Genomics’ protocols (CG000375 Rev B for tissue, CG000124 Rev E for organoids). Briefly, snap-frozen tissue was minced into smaller fragments using a scalpel and then added to lysis buffer (Thermo Fisher Scientific, Cat#PI28324). Tissue was homogenized using a pellet pestle 15 times then incubated for 5 minutes in lysis buffer (1 minute if organoids). The cell suspension was filtered through a 33 μm strainer. Suspension was centrifuged at 500g for 5 mins at 4°C. Supernatant was removed and the pellet was washed with PBS + 1% BSA, centrifuged at 500g for 5 mins at 4°C; pellet was washed and centrifuged again. Permeabilization for ATAC-seq samples was performed by incubating the pellet in 0.1X lysis buffer and incubated for 2 minutes. Suspension was then centrifuged at 500g for 5 mins at 4°C; supernatant was removed and pellet was resuspended in diluted nuclei buffer.

### Single Nuclei Analysis Overview

Data matrices for further analysis were generated using the CellRanger pipeline (v) under standard parameters by the University of Michigan Advanced Genomics Sequencing Core. Cells were re-called on raw matrices using Cellbender^42^ v0.3, which also corrected for ambient RNA at an FPR of 0.01. ATAC-sequencing data was not corrected. Corrected gene expression matrices were important into Seurat^43^ v4 and un-corrected ATAC-sequencing were imported into Signac^44^ v1 in RStudio v1.4 running on R 4.4 for further analysis. A distal sample from HT560 was excluded from further analysis due to very low feature complexity.

### Single Nuclei Gene Expression

#### Preprocessing/QC Filtering (All Data)

For analysis of gene expression data only, cells were eliminated that did not fit the following parameters: between 1000 to 5000 features, less than 5% and mitochondria RNA and 7.5% ribosomal RNA reads and greater than 1000 unique molecular identifiers (UMIs). All 15 gene expression datasets underwent independent normalization, variable feature selection, scaling and principal component analysis (PCA). Anchors for batch-corrected integration were determined using a reciprocal PCA approach considering 30 principle components (PCs) from each dataset. Integrated data matrices were produced, scaled, subjected to PCA, which formed the basis for UMAP projections (30 PCs), cell-neighbor identification (30 PCs) and cell clustering (resolution = 0.75).

#### Data Integration, Dimensional Reduction, Clustering and Additional QC (Tissue Data)

All 15 gene expression datasets underwent independent normalization, variable feature selection, scaling and principal component analysis (PCA). Anchors for batch-corrected integration were determined using a reciprocal PCA approach considering 30 principle components (PCs) from each dataset. Integrated data matrices were produced, scaled, subjected to PCA, which formed the basis for UMAP projection (30 PCs), cell-neighbor identification (30 PCs) and cell clustering (resolution = 0.75). At this stage a cluster of cells in the mesenchyme defined by higher mitochondrial and ribosomal content than all other clusters was removed, as well as a cluster of cells in the Epithelium that was contaminated with mesenchymal marker expression. After removal of these low-confidence cell clusters data was freshly integrated by reciprocal PCA, dimensionally reduced and clustered as above.

#### Data Integration, Dimensional Reduction, Clustering and Additional QC (Organoid Data)

4 gene expression datasets from positional organoids were normalized and underwent variable feature selection, scaling and principal component analysis (PCA) independently. Anchors for batch-corrected integration were determined using a reciprocal PCA approach considering 30 PCs from each dataset. Integrated data matrices were produced, scaled, subjected to PCA, which formed the basis for UMAP projection (16 PCs), cell-neighbor identification (16 PCs) and cell clustering (resolution = 0.5). A small cluster of mesenchyme-like cells was removed at this stage and integration, PCA and UMAP was performed as above, this time utilizing 20 PCs for UMAP and neighbor finding and 0.5 resolution for determining clusters.

#### Referenced-based mapping (label transfer) of positional organoids

To classify cell types in positional organoid data according to their most transcriptionally similar in vivo counterparts we performed reference-based mapping in Seurat using organoid data as query and the PCA data from tissue as reference. Transfer anchors were first identified by FindTransferAnchors() using common features to the integrated assays of both datasets, the PCA structure of the reference dataset and setting k.filter = 200. MapQuery() was used to transfer labels using level 3 annotations and embed query cells onto the reference UMAP.

#### Cell Type Annotation

Major classes of cell types in the lung (Epithelial, Mesenchymal, Immune, Endothelial and Neuronal) were first annotated using canonical markers that distinguish each major class (Figure S1C). Sub clustering of each major cell class was performed using the integrated data matrix from reciprocal PCA integration to generate a new PCA for each cell class. Data was displayed by UMAP. Cell neighbors were calculated and cell clustering was performed using cell class specific parameters (Epithelium: 15 PCs, 0.6 resolution; Mesenchyme: 18 PCs, 0.8 resolution; Neuronal: 7 PCs, 0.2 resolution, Endothelial 5 PCs, 0.125 resolution; Immune 10 PCs, 0.15 resolution). Additional sub clustering using the above workflow was performed on large clusters of cells within the epithelium and immune compartments (Basal Cells 6 PCs, 0.3 resolution; Distal Epithelial Cells: 9 PCs, 0.5 resolution, Lymphoid 5 PCs, 0.15 resolution). Gene enrichment lists were generated using the FindMarkers() function to run Wilcoxon Ranked Sum test, filtering out mitochondrial, ribosomal, genes with less than 25% expression and less than 0.25 log-fold change within the cluster of interest. Using highly enriched canonical markers of known cell types we annotated cells in our gene expression data at multiple levels of coarse and fine annotation (Supplemental Table 3), for example KRT4+ basal cells (Level 1:Epithelial, Level 2:Basal, Level 3: KRT4+ basal) or mature chondrocytes (Level 1: Mesenchymal, Level 2: Chondrocyte Lineage, Level 3: Mature Chondrocyte).

### Xenium Data Analysis

We followed the standard Seurat pipeline to load Xenium spatial transcriptomic data in R. We applied log normalization using the NormalizeData function onto Xenium data. snRNA-seq data from human fetal lung micro-dissected regions was normalized using the same function after it was subsetted with the same genes in the Xenium data set. Then, we performed Seurat recommended pipeline to transfer labels from snRNA-seq data (reference) to Xenium spatial transcriptomic data (query). Anchors between reference and query object were found by the function FindTransferAnchors with the default setting. Function TransferData was then applied to transfer labels from the reference to query object. After adding the predicted labels calculated by TransferData to Xenium data set, we were able to export cell ID with the transferred labels as csv file, using which as the input to Xenium Explorer V3.1.0. Data was visualized in Xenium Explorer using built-in functions.

## ACKNOWLEDGMENTS

We would like to thank members of the Spence lab for their helpful discussions and technical support. We would also like to thank the University of Michigan Advanced Genomics Core and the University of Washington Laboratory of Developmental Biology. We express gratitude to the patients who donated tissue samples to make this research possible. This work was supported in part by the Chan Zuckerberg Initiative (CZI) Seed Network (2019-002440) and CZI Pediatric Network (2021-237566), an advised fund of the Silicon Valley Community Foundation, and also the National Heart, Lung, and Blood Institute (NHLBI; R01HL119215 and R01HL166139) to J.R.S.; and the Judith Tam ALK Lung Cancer Research Initiative to P.P.H. and S.D.M. P.P.H. received support from the Rogel Cancer Center Fellowship. S.D.M. is supported by the Breast Cancer Research Foundation. A.S.C. received support from a T32 Michigan Medical Scientist Training Program (5T32GM007863-40) and a Ruth L. Kirschstein Predoctoral Individual National Research Service Award (NIH-NHLBI F30HL156474). T.F. received support from a NIH Tissue Engineering and Regenerative Medicine Training Grant (NIH-NIDCR T32DE007057).

---

**Supplemental Figure 1.**
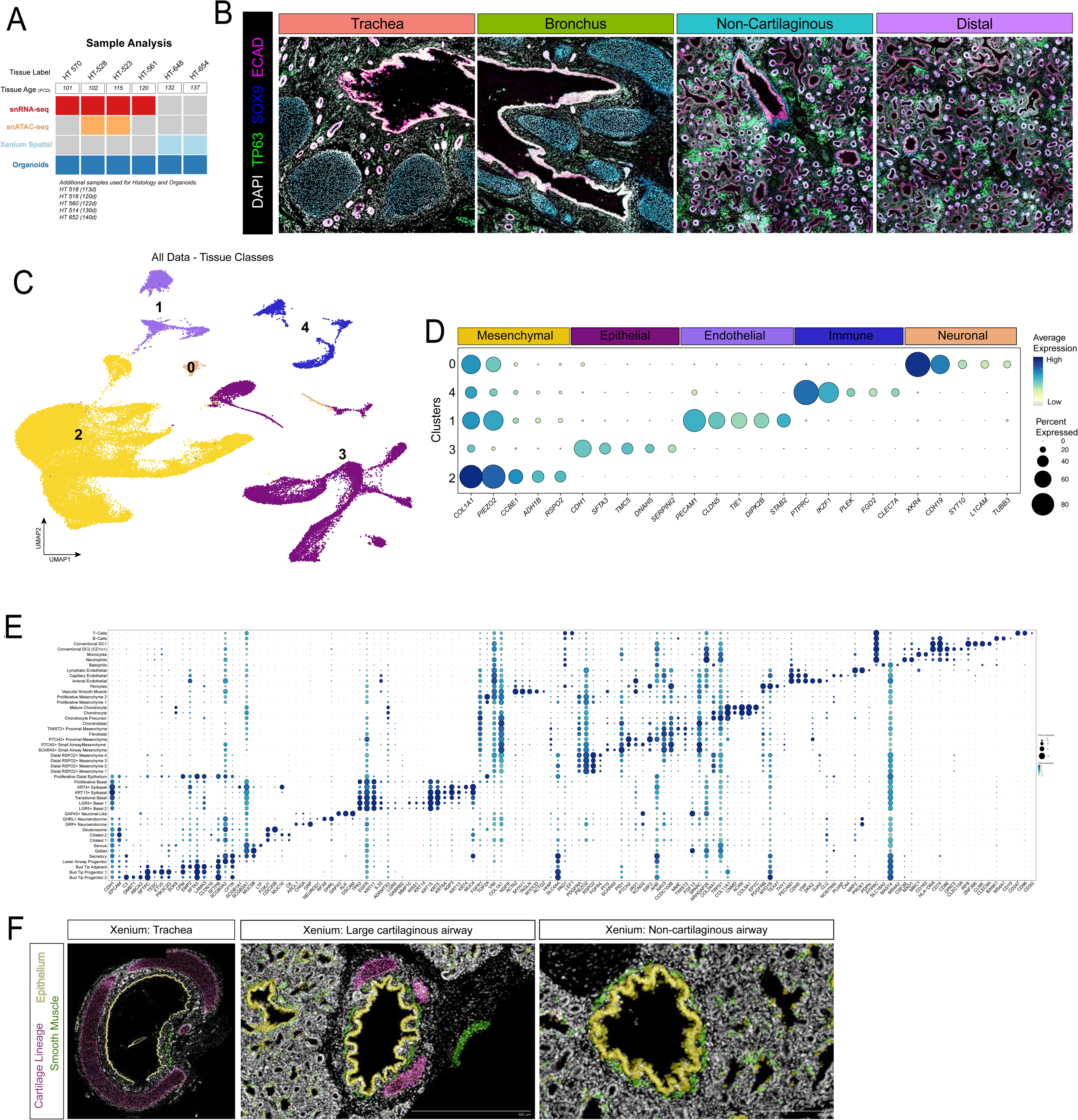
A. Schematic highlighting specific airway regions that were dissected and summary of analysis performed across all 7 tissue samples with all 4 regions including snRNA-seq, snATAC-seq, Xenium spatial transcriptomics, and derived organoid lines from each region. B. Representative immunofluorescence (IF) image of 113 PCD lung for regions of interest: trachea, bronchus, non-cartilaginous airway and distal lung for canonical cell and tissue type markers including TP63, ECAD, FOXJ1, SOX9. C. UMAP cluster plot of snRNA-seq from human fetal lung micro-dissected regions, color coded by tissue compartment. D. Dot plot for expression of top genes defining each tissue compartment cluster of UMAP in (S1C). E. Dot plot for expression of top 3 genes defining each cluster of UMAP in Figure 1A. F. Xenium spatial transcriptomic images of trachea, cartilaginous airways, and non-cartilaginous airways with epithelium (yellow), smooth muscle (green) and cartilage (pink) highlighted.

**Supplemental Figure 2.**
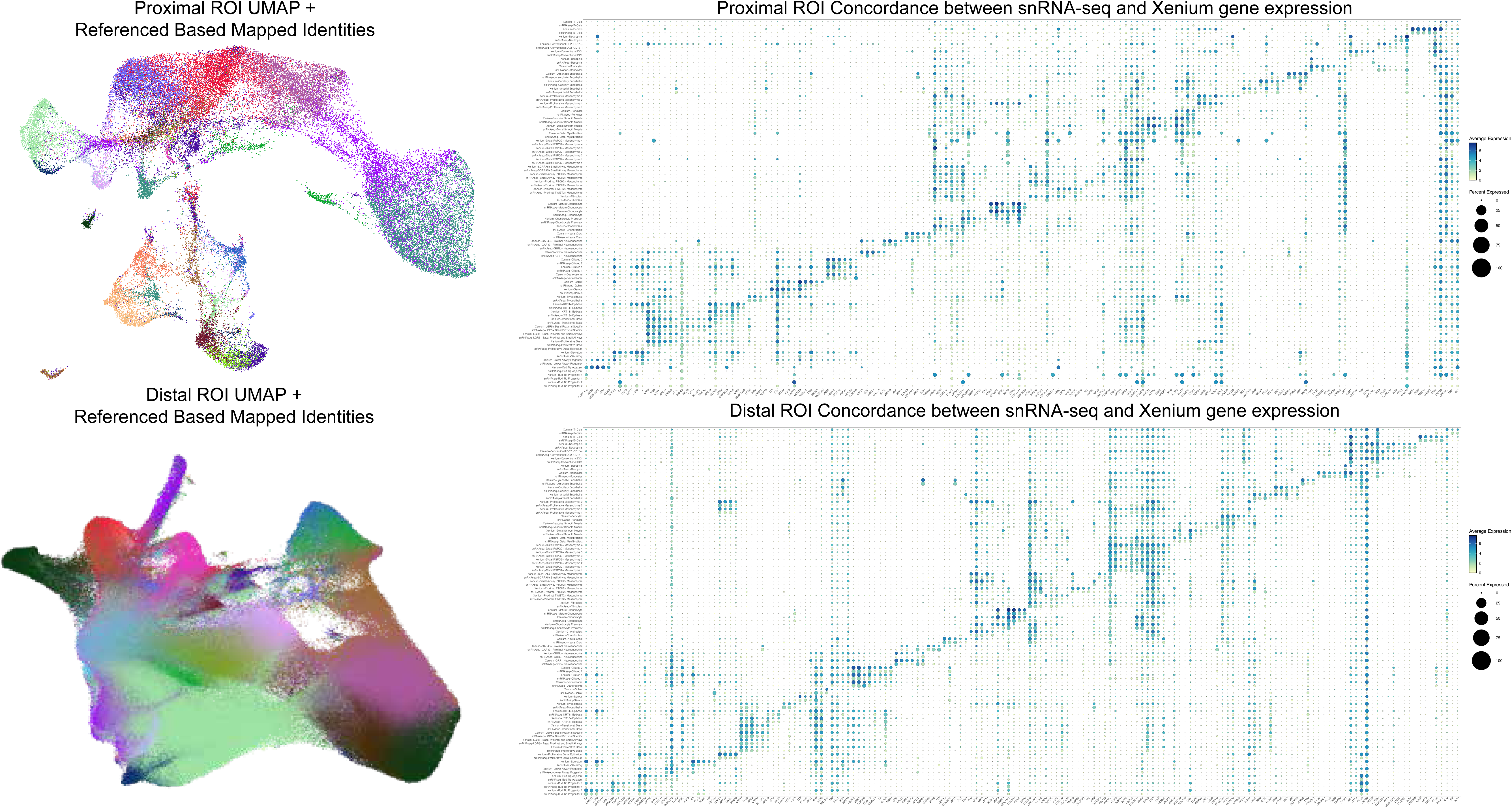
UMAP cluster plot and dotplots correlating label transferred identities of snRNA-seq data onto Xenium spatial transcriptomic datasets of trachea (top) and lung (bottom) samples.

**Supplemental Figure 3.**
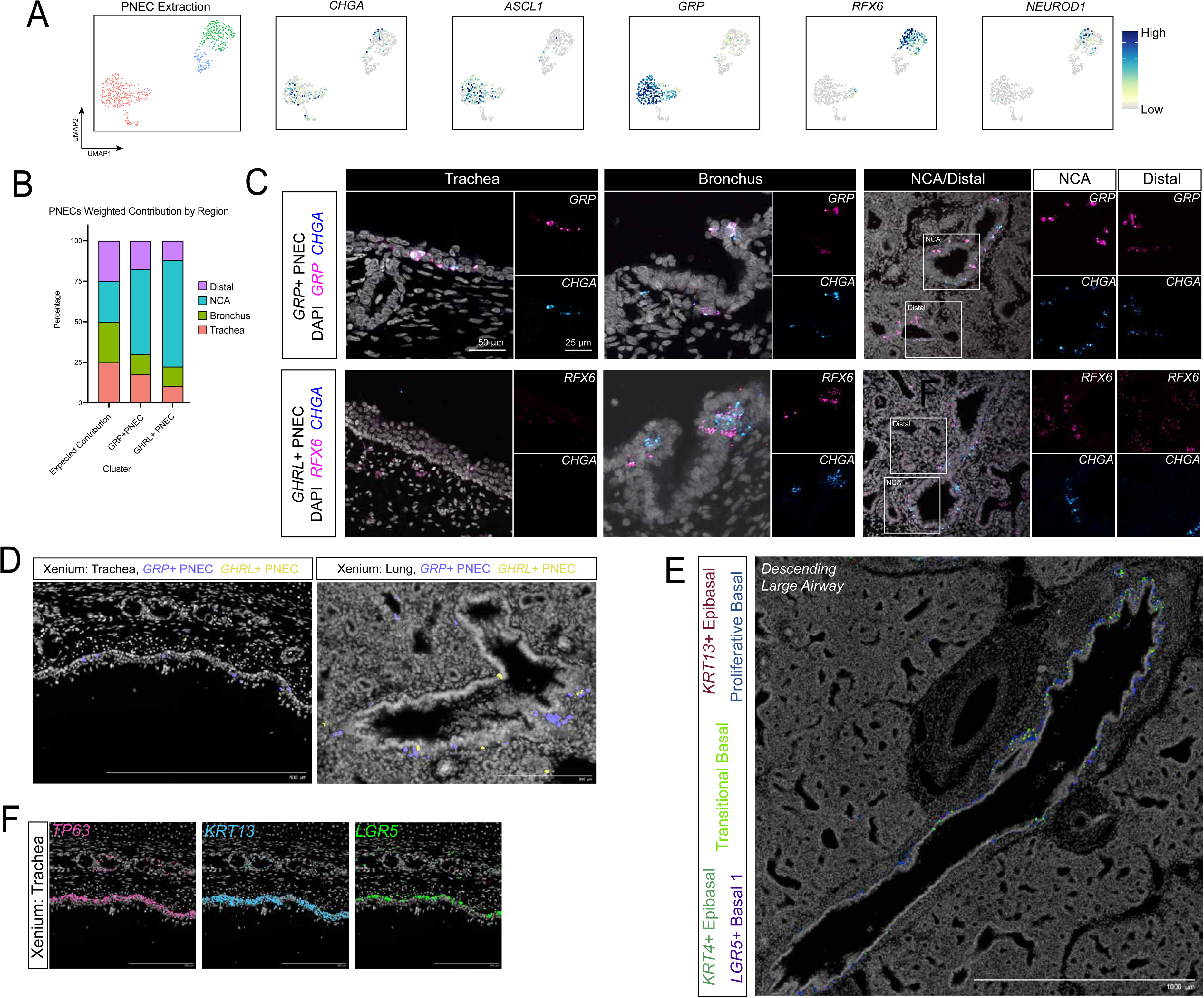
A. UMAP cluster plot for computationally extracted pulmonary neuroendocrine cells (PNECs), demonstrating 2 distinct clusters and PNEC sub-types. Feature plots for canonical PNEC markers (*ASCL1, CHGA*) and sub-type defining genes (*GRP*, *RFX6*, *NEUROD1*). B. Distribution plot quantifies weighted percent of each PNEC sub-type across 4 regions. C. Representative FISH images with co-IF for PNEC sub-types across 4 distinct airway regions. D. Xenium spatial transcriptomics image of 132d human fetal trachea (left) and lung (right) for *GRP*+ PNECs and *GHRL*+ PNECs identified by label transfer from snRNA-seq in Figure 1A. E. Xenium spatial transcriptomics image of basal cell sub-types in descending large airway. F. Xenium spatial transcriptomic images of *TP63*, *KRT13* and *LGR5* transcript expression in 132d human fetal trachea.

**Supplemental Figure 4.**
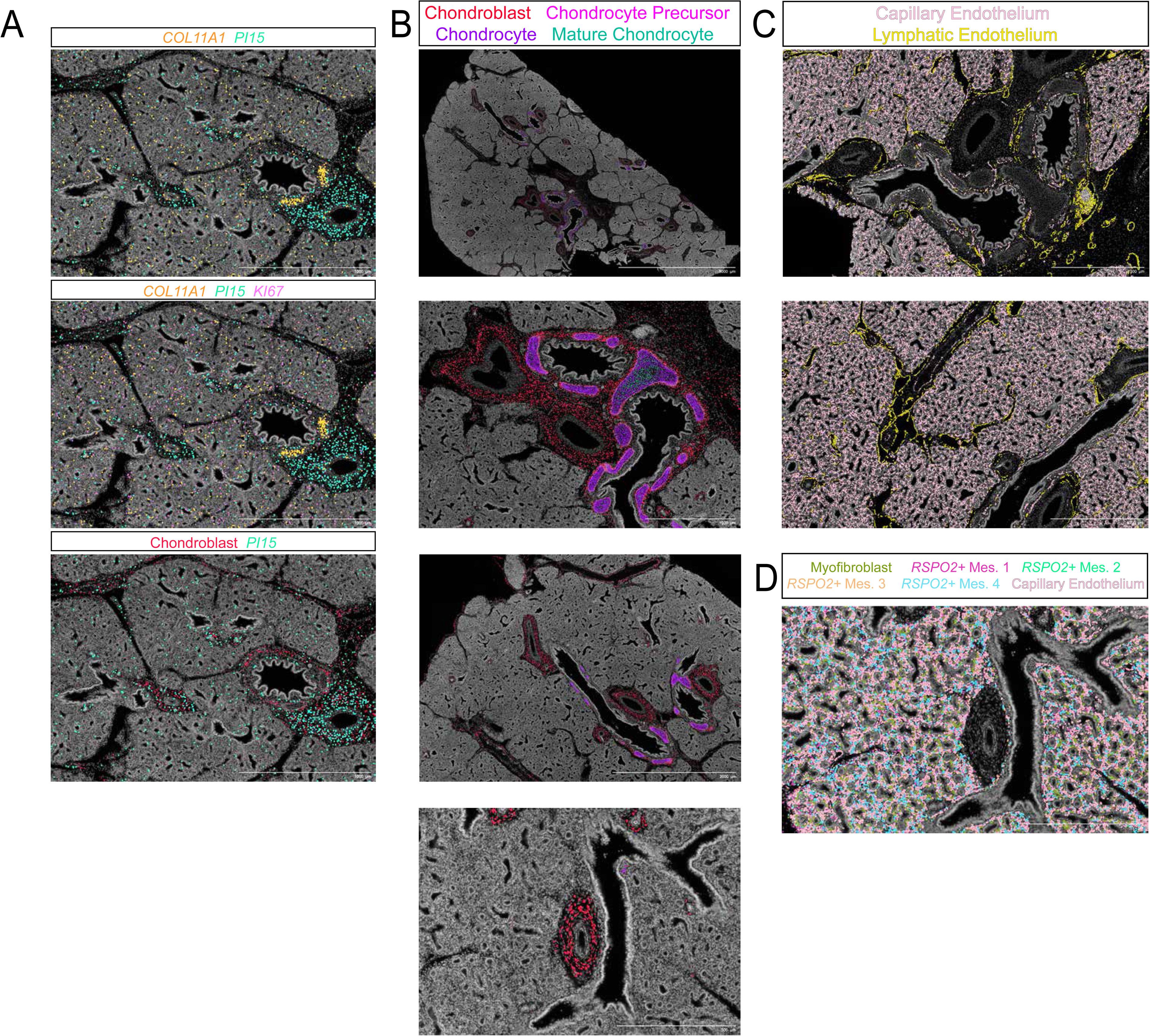
A. Xenium spatial transcriptomic images of 132d human fetal lung *COL11A1*, *PI15,* and *KI67* transcript expression in lower, cartilaginous and non-cartilaginous airways. Bottom image includes Chondroblast cells identified by label transfer. B. Xenium spatial transcriptomics image of 132d human fetal lung for Chondroblast, Chondrocyte Precursor, Chondrocytes, and Mature Chondrocytes color coded by label transfer from identities defined in snRNA dataset (Figure 1A). Top image, entire lung section; top middle, bronchus; bottom middle, lower cartilaginous airways; bottom, non- cartilaginous and distal airways. C. Xenium spatial transcriptomics image of 132d human fetal lung for Capillary and Lymphatic endothelium in bronchus (top) and lower airways (bottom). D. Xenium spatial transcriptomics image of 132d human fetal lung for distal mesenchymal subtypes alongside Capillary endothelium.

**Supplemental Figure 5.**
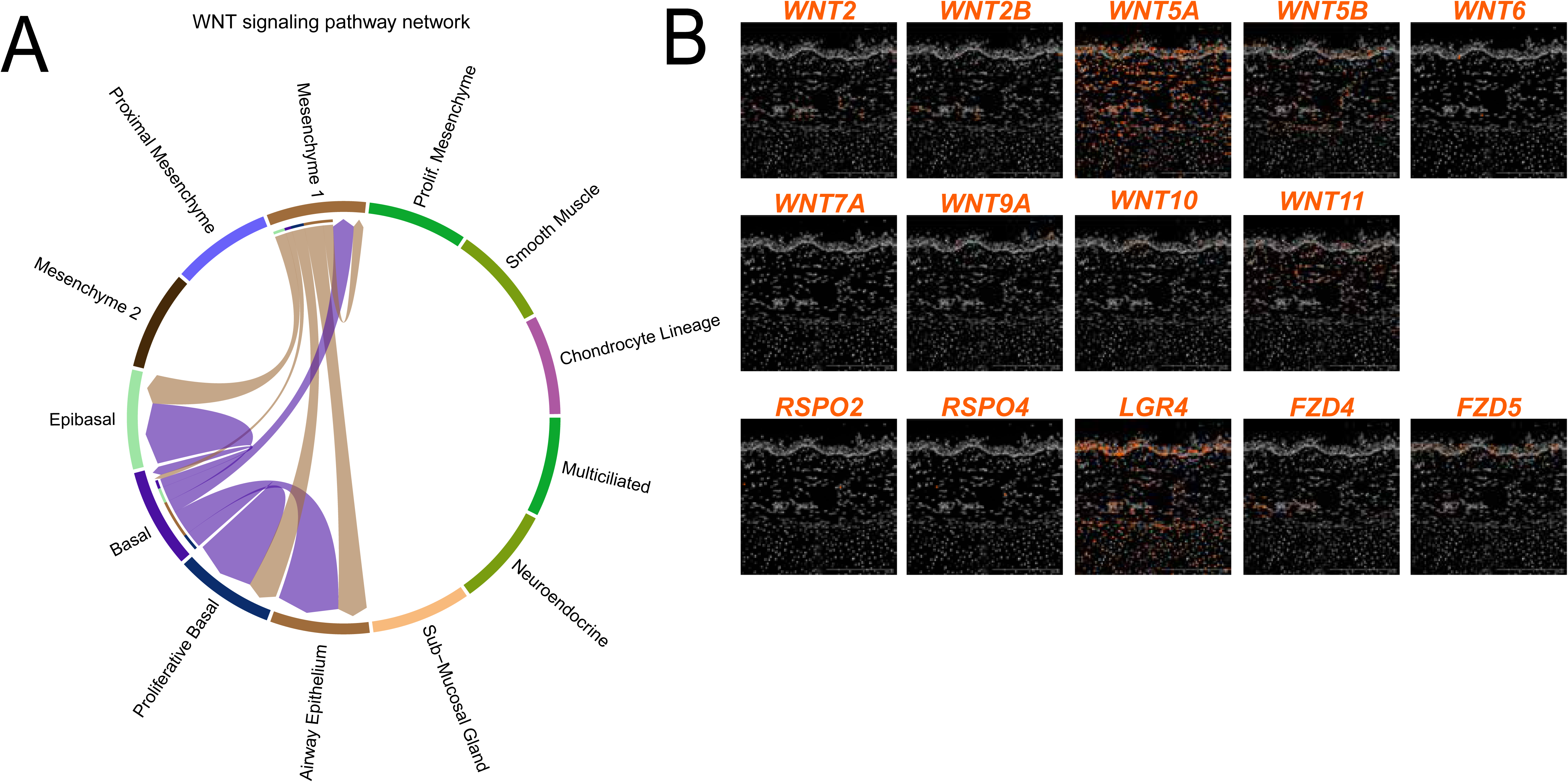
A. Chord plot for CellChat analysis of canonical WNT signaling components on (A) clusters. B. Xenium spatial transcriptomics image of WNT signaling components transcript expression in 132d human fetal trachea.

**Supplemental Figure 6.**
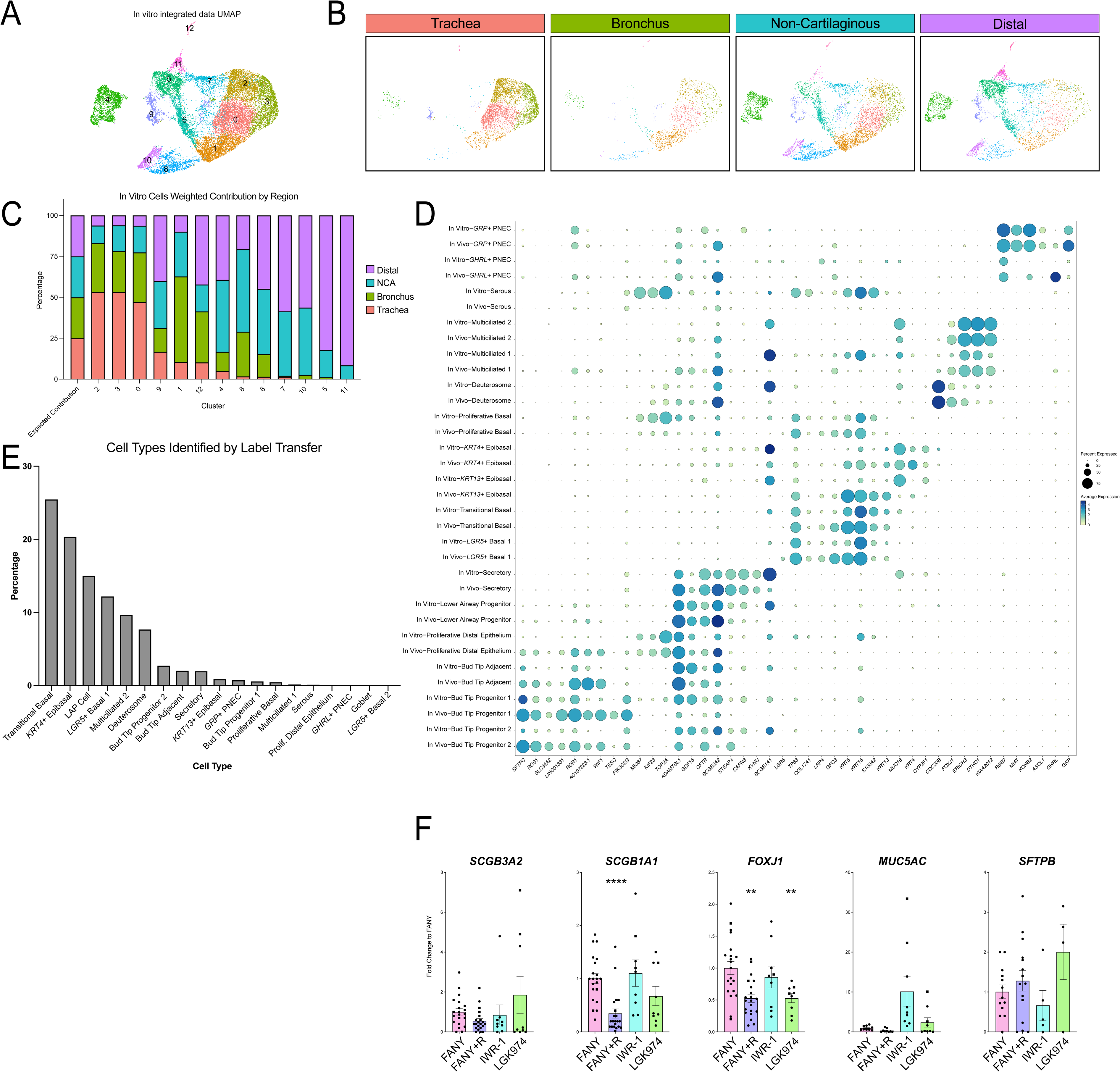
A. UMAP cluster plot of snRNA-seq data from organoid lines (n=1) combining all 4 regions. B. UMAP cluster plot from (A) split by regional airway organoid line. C. Distribution plot quantifies weighted percent of clusters derived from each regional organoid line. D. Dot plot for expression of defining cluster genes correlating in vivo and in vitro clusters. E. Percentage of cells in organoid snRNA-seq as identified by label transfer from in vivo snRNA-seq data from Figure 2A. F. RT-qPCR of airway cell markers in WNT modulation experiment. This quantification was performed on a minimum of three separate biological replicates with three technical replicates each. Error bars represent standard error of the mean. Statistical tests were performed by Welch ANOVA test with Dunnett’s T3 multiple comparisons. P-values ** < 0.005, *** < 0.0005, **** < 0.0001.

## REFERENCES

1. Zepp, J.A., and Morrisey, E.E. (2019). Cellular crosstalk in the development and regeneration of the respiratory system. Nat. Rev. Mol. Cell Biol. 20, 551–566. 10.1038/s41580-019-0141-3.

2. Conway, R.F., Frum, T., Conchola, A.S., and Spence, J.R. (2020). Understanding Human Lung Development through In Vitro Model Systems. BioEssays 42, 2000006. 10.1002/bies.202000006.

3. Nikolić, M.Z., Caritg, O., Jeng, Q., Johnson, J.-A., Sun, D., Howell, K.J., Brady, J.L., Laresgoiti, U., Allen, G., Butler, R., et al. (2017). Human embryonic lung epithelial tips are multipotent progenitors that can be expanded in vitro as long-term self-renewing organoids. eLife. 10.7554/eLife.26575.

4. Arason, A.J., Jonsdottir, H.R., Halldorsson, S., Benediktsdottir, B.E., Bergthorsson, J.T., Ingthorsson, S., Baldursson, O., Sinha, S., Gudjonsson, T., and Magnusson, M.K. (2014). deltaNp63 Has a Role in Maintaining Epithelial Integrity in Airway Epithelium. PLOS ONE 9, e88683. 10.1371/journal.pone.0088683.

5. Daniely, Y., Liao, G., Dixon, D., Linnoila, R.I., Lori, A., Randell, S.H., Oren, M., and Jetten, A.M. (2004). Critical role of p63 in the development of a normal esophageal and tracheobronchial epithelium. Am. J. Physiol.-Cell Physiol. 287, C171–C181. 10.1152/ajpcell.00226.2003.

6. Miller, A.J., Yu, Q., Czerwinski, M., Tsai, Y.-H., Conway, R.F., Wu, A., Holloway, E.M., Walker, T., Glass, I.A., Treutlein, B., et al. (2020). In Vitro and In Vivo Development of the Human Airway at Single-Cell Resolution. Dev. Cell 53, 117–128.e6. 10.1016/j.devcel.2020.01.033.

7. Sountoulidis, A., Marco Salas, S., Braun, E., Avenel, C., Bergenstråhle, J., Theelke, J., Vicari, M., Czarnewski, P., Liontos, A., Abalo, X., et al. (2023). A topographic atlas defines developmental origins of cell heterogeneity in the human embryonic lung. Nat. Cell Biol. 25, 351–365. 10.1038/s41556-022-01064-x.

8. He, P., Lim, K., Sun, D., Pett, J.P., Jeng, Q., Polanski, K., Dong, Z., Bolt, L., Richardson, L., Mamanova, L., et al. (2022). A human fetal lung cell atlas uncovers proximal-distal gradients of differentiation and key regulators of epithelial fates. Cell 185, 4841–4860.e25. 10.1016/j.cell.2022.11.005.

9. Danopoulos, S., Alonso, I., Thornton, M.E., Grubbs, B.H., Bellusci, S., Warburton, D., and Al Alam, D. (2018). Human lung branching morphogenesis is orchestrated by the spatiotemporal distribution of ACTA2, SOX2, and SOX9. Am. J. Physiol.-Lung Cell. Mol. Physiol. 314, L144–L149. 10.1152/ajplung.00379.2017.

10. Trecartin, A., Danopoulos, S., Spurrier, R., Knaneh-Monem, H., Hiatt, M., Driscoll, B., Hochstim, C., Al-Alam, D., and Grikscheit, T.C. (2016). Establishing Proximal and Distal Regional Identities in Murine and Human Tissue-Engineered Lung and Trachea. Tissue Eng. Part C Methods 22, 1049–1057. 10.1089/ten.tec.2016.0261.

11. Quach, H., Farrell, S., Wu, M.J.M., Kanagarajah, K., Leung, J.W.-H., Xu, X., Kallurkar, P., Turinsky, A.L., Bear, C.E., Ratjen, F., et al. (2024). Early human fetal lung atlas reveals the temporal dynamics of epithelial cell plasticity. Nat. Commun. 15, 5898. 10.1038/s41467-024-50281-5.

12. Conchola, A.S., Frum, T., Xiao, Z., Hsu, P.P., Kaur, K., Downey, M.S., Hein, R.F.C., Miller, A.J., Tsai, Y.-H., Wu, A., et al. (2023). Regionally distinct progenitor cells in the lower airway give rise to neuroendocrine and multiciliated cells in the developing human lung. Proc. Natl. Acad. Sci. 120, e2210113120. 10.1073/pnas.2210113120.

13. Zhou, Y., Yang, Y., Guo, L., Qian, J., Ge, J., Sinner, D., Ding, H., Califano, A., and Cardoso, W.V. (2022). Airway basal cells show regionally distinct potential to undergo metaplastic differentiation. eLife 11, e80083. 10.7554/eLife.80083.

14. Lin, B., Shah, V.S., Chernoff, C., Sun, J., Shipkovenska, G.G., Vinarsky, V., Waghray, A., Xu, J., Leduc, A.D., Hintschich, C.A., et al. (2024). Airway hillocks are injury-resistant reservoirs of unique plastic stem cells. Nature 629, 869–877. 10.1038/s41586-024-07377-1.

15. Montoro, D.T., Haber, A.L., Biton, M., Vinarsky, V., Lin, B., Birket, S.E., Yuan, F., Chen, S., Leung, H.M., Villoria, J., et al. (2018). A revised airway epithelial hierarchy includes CFTR-expressing ionocytes. Nature 560, 319–324. 10.1038/s41586-018-0393-7.

16. Rim, E.Y., Clevers, H., and Nusse, R. (2022). The Wnt Pathway: From Signaling Mechanisms to Synthetic Modulators. Annu. Rev. Biochem. 91, 571–598. 10.1146/annurev-biochem-040320-103615.

17. Hein, R.F.C., Wu, J.H., Holloway, E.M., Frum, T., Conchola, A.S., Tsai, Y.-H., Wu, A., Fine, A.S., Miller, A.J., Szenker-Ravi, E., et al. (2022). R-SPONDIN2+ mesenchymal cells form the bud tip progenitor niche during human lung development. Dev. Cell 57, 1598–1614.e8. 10.1016/j.devcel.2022.05.010.

18. Jacob, A., Morley, M., Hawkins, F., McCauley, K.B., Jean, J.C., Heins, H., Na, C.-L., Weaver, T.E., Vedaie, M., Hurley, K., et al. (2017). Differentiation of Human Pluripotent Stem Cells into Functional Lung Alveolar Epithelial Cells. Cell Stem Cell 21, 472–488.e10. 10.1016/j.stem.2017.08.014.

19. Lim, K., Donovan, A.P.A., Tang, W., Sun, D., He, P., Pett, J.P., Teichmann, S.A., Marioni, J.C., Meyer, K.B., Brand, A.H., et al. (2023). Organoid modeling of human fetal lung alveolar development reveals mechanisms of cell fate patterning and neonatal respiratory disease. Cell Stem Cell 30, 20–37.e9. 10.1016/j.stem.2022.11.013.

20. Frum, T., Hsu, P.P., Hein, R.F.C., Conchola, A.S., Zhang, C.J., Utter, O.R., Anand, A., Zhang, Y., Clark, S.G., Glass, I., et al. (2023). Opposing roles for TGFβ- and BMP-signaling during nascent alveolar differentiation in the developing human lung. Npj Regen. Med. 8, 1–19. 10.1038/s41536-023-00325-z.

21. Luca, V.C., Miao, Y., Li, X., Hollander, M.J., Kuo, C.J., and Garcia, K.C. (2020). Surrogate R-spondins for tissue-specific potentiation of Wnt Signaling. PLOS ONE 15, e0226928. 10.1371/journal.pone.0226928.

22. Carmon, K.S., Gong, X., Lin, Q., Thomas, A., and Liu, Q. (2011). R-spondins function as ligands of the orphan receptors LGR4 and LGR5 to regulate Wnt/β-catenin signaling. Proc. Natl. Acad. Sci. 108, 11452–11457. 10.1073/pnas.1106083108.

23. de Lau, W., Barker, N., Low, T.Y., Koo, B.-K., Li, V.S.W., Teunissen, H., Kujala, P., Haegebarth, A., Peters, P.J., van de Wetering, M., et al. (2011). Lgr5 homologues associate with Wnt receptors and mediate R-spondin signalling. Nature 476, 293–297. 10.1038/nature10337.

24. Hou, Z., Wu, Q., Sun, X., Chen, H., Li, Y., Zhang, Y., Mori, M., Yang, Y., Que, J., and Jiang, M. (2019). Wnt/Fgf crosstalk is required for the specification of basal cells in the mouse trachea. Development 146, dev171496. 10.1242/dev.171496.

25. Deprez, M., Zaragosi, L.-E., Truchi, M., Becavin, C., Ruiz García, S., Arguel, M.-J., Plaisant, M., Magnone, V., Lebrigand, K., Abelanet, S., et al. (2020). A Single-Cell Atlas of the Human Healthy Airways. Am. J. Respir. Crit. Care Med. 202, 1636–1645. 10.1164/rccm.201911-2199OC.

26. Jin, S., Guerrero-Juarez, C.F., Zhang, L., Chang, I., Ramos, R., Kuan, C.-H., Myung, P., Plikus, M.V., and Nie, Q. (2021). Inference and analysis of cell-cell communication using CellChat. Nat. Commun. 12, 1088. 10.1038/s41467-021-21246-9.

27. Brechbuhl, H.M., Ghosh, M., Smith, M.K., Smith, R.W., Li, B., Hicks, D.A., Cole, B.B., Reynolds, P.R., and Reynolds, S.D. (2011). β-catenin dosage is a critical determinant of tracheal basal cell fate determination. Am. J. Pathol. 179, 367–379. 10.1016/j.ajpath.2011.03.016.

28. Ishii, Y., Orr, J.C., El Mdawar, M.-B., de Pilger, D.R.B., Pearce, D.R., Lazarus, K.A., Graham, R.E., Nikolić, M.Z., Ketteler, R., Carragher, N.O., et al. (2024). Compound screening in primary human airway basal cells identifies Wnt pathway activators as potential pro-regenerative therapies. bioRxiv, 2024.08.13.606573. 10.1101/2024.08.13.606573.

29. Aros, C.J., Vijayaraj, P., Pantoja, C.J., Bisht, B., Meneses, L.K., Sandlin, J.M., Tse, J.A., Chen, M.W., Purkayastha, A., Shia, D.W., et al. (2020). Distinct Spatiotemporally Dynamic Wnt-Secreting Niches Regulate Proximal Airway Regeneration and Aging. Cell Stem Cell 27, 413–429.e4. 10.1016/j.stem.2020.06.019.

30. Haas, M., Vázquez, J.L.G., Sun, D.I., Tran, H.T., Brislinger, M., Tasca, A., Shomroni, O., Vleminckx, K., and Walentek, P. (2019). ΔN-Tp63 Mediates Wnt/β-Catenin-Induced Inhibition of Differentiation in Basal Stem Cells of Mucociliary Epithelia. Cell Rep. 28, 3338–3352.e6. 10.1016/j.celrep.2019.08.063.

31. Zhang, Y., Black, K.E., Phung, T.-K.N., Thundivalappil, S.R., Lin, T., Wang, W., Xu, J., Zhang, C., Hariri, L.P., Lapey, A., et al. (2023). Human Airway Basal Cells Undergo Reversible Squamous Differentiation and Reshape Innate Immunity. Am. J. Respir. Cell Mol. Biol. 68, 664–678. 10.1165/rcmb.2022-0299OC.

32. Polkoff, K.M., Lampe, R., Gupta, N.K., Murphy, Y., Chung, J., Carter, A., Simon, J.M., Gleason, K., Moatti, A., Murthy, P.K., et al. (2024). Novel porcine model reveals two distinct LGR5 cell types during lung development and homeostasis. bioRxiv, 2022.12.09.516617. 10.1101/2022.12.09.516617.

33. Hill, A.B.T., Murphy, Y.M., Polkoff, K.M., Edwards, L., Walker, D.M., Moatti, A., Greenbaum, A., and Piedrahita, J.A. (2024). A gene edited pig model for studying LGR5(+) stem cells: implications for future applications in tissue regeneration and biomedical research. Front. Genome Ed. 6, 1401163. 10.3389/fgeed.2024.1401163.

34. Lee, J.-H., Tammela, T., Hofree, M., Choi, J., Marjanovic, N.D., Han, S., Canner, D., Wu, K., Paschini, M., Bhang, D.H., et al. (2017). Anatomically and Functionally Distinct Lung Mesenchymal Populations Marked by Lgr5 and Lgr6. Cell 170, 1149–1163.e12. 10.1016/j.cell.2017.07.028.

35. King, N.E., Suzuki, S., Barillà, C., Hawkins, F.J., Randell, S.H., Reynolds, S.D., Stripp, B.R., and Davis, B.R. (2020). Correction of Airway Stem Cells: Genome Editing Approaches for the Treatment of Cystic Fibrosis. Hum. Gene Ther. 31, 956–972. 10.1089/hum.2020.160.

36. Carraro, G., Mulay, A., Yao, C., Mizuno, T., Konda, B., Petrov, M., Lafkas, D., Arron, J.R., Hogaboam, C.M., Chen, P., et al. (2020). Single-Cell Reconstruction of Human Basal Cell Diversity in Normal and Idiopathic Pulmonary Fibrosis Lungs. Am. J. Respir. Crit. Care Med. 202, 1540–1550. 10.1164/rccm.201904-0792OC.

37. Aros, C.J., Pantoja, C.J., and Gomperts, B.N. (2021). Wnt signaling in lung development, regeneration, and disease progression. Commun. Biol. 4, 601. 10.1038/s42003-021-02118-w.

38. Lynch, T.J., Anderson, P.J., Xie, W., Crooke, A.K., Liu, X., Tyler, S.R., Luo, M., Kusner, D.M., Zhang, Y., Neff, T., et al. (2016). Wnt Signaling Regulates Airway Epithelial Stem Cells in Adult Murine Submucosal Glands. Stem Cells Dayt. Ohio 34, 2758–2771. 10.1002/stem.2443.

39. Hsu, H.-S., Liu, C.-C., Lin, J.-H., Hsu, T.-W., Su, K., and Hung, S.-C. (2014). Repair of naphthalene-induced acute tracheal injury by basal cells depends on β-catenin. J. Thorac. Cardiovasc. Surg. 148, 322–332. 10.1016/j.jtcvs.2013.10.039.

40. Giangreco, A., Lu, L., Vickers, C., Teixeira, V.H., Groot, K.R., Butler, C.R., Ilieva, E.V., George, P.J., Nicholson, A.G., Sage, E.K., et al. (2012). β-Catenin determines upper airway progenitor cell fate and preinvasive squamous lung cancer progression by modulating epithelial-mesenchymal transition. J. Pathol. 226, 575–587. 10.1002/path.3962.

41. Childs, C.J., Holloway, E.M., Sweet, C.W., Tsai, Y.-H., Wu, A., Vallie, A., Eiken, M.K., Capeling, M.M., Zwick, R.K., Palikuqi, B., et al. (2023). EPIREGULIN creates a developmental niche for spatially organized human intestinal enteroids. JCI Insight 8. 10.1172/jci.insight.165566.

42. Fleming, S.J., Chaffin, M.D., Arduini, A., Akkad, A.-D., Banks, E., Marioni, J.C., Philippakis, A.A., Ellinor, P.T., and Babadi, M. (2023). Unsupervised removal of systematic background noise from droplet-based single-cell experiments using CellBender. Nat. Methods 20, 1323–1335. 10.1038/s41592-023-01943-7.

43. Stuart, T., Butler, A., Hoffman, P., Hafemeister, C., Papalexi, E., Mauck, W.M., Hao, Y., Stoeckius, M., Smibert, P., and Satija, R. (2019). Comprehensive Integration of Single-Cell Data. Cell 177, 1888–1902.e21. 10.1016/j.cell.2019.05.031.

44. Stuart, T., Srivastava, A., Madad, S., Lareau, C.A., and Satija, R. (2021). Single-cell chromatin state analysis with Signac. Nat. Methods 18, 1333–1341. 10.1038/s41592-021-01282-5.

45. Zhang, Y., Liu, T., Meyer, C.A., Eeckhoute, J., Johnson, D.S., Bernstein, B.E., Nusbaum, C., Myers, R.M., Brown, M., Li, W., et al. (2008). Model-based Analysis of ChIP-Seq (MACS). Genome Biol. 9, R137. 10.1186/gb-2008-9-9-r137.

46. Cusanovich, D.A., Daza, R., Adey, A., Pliner, H.A., Christiansen, L., Gunderson, K.L., Steemers, F.J., Trapnell, C., and Shendure, J. (2015). Multiplex single-cell profiling of chromatin accessibility by combinatorial cellular indexing. Science 348, 910–914. 10.1126/science.aab1601.

