## Supplemental Table 1 for "A spatially-resolved blueprint of the developing human lung reveals a WNT-driven niche for basal stem cells"

| **T-Cells** | **Monocytes** | **B-Cells** | **CD2+** | **DC1** | **Neutrophils** | **Basophils** |
| --- | --- | --- | --- | --- | --- | --- |
| BCL11B | F13A1 | EBF1 | HLA-DRA | CCSER1 | VCAN | RAB27B |
| PRKCH | CD163L1 | BANK1 | HLA-DPB1 | WDFY4 | ACSL1 | DNM3 |
| CD247 | CD36 | PAX5 | ENTPD1-AS1 | CLNK | THBS1 | SLC18A2 |
| SKAP1 | MRC1 | NIBAN3 | HLA-DRB1 | CADM1 | NAMPT | LINC02284 |
| THEMIS | DAB2 | RALGPS2 | CD86 | CPVL | FCN1 | LTBP1 |
| CAMK4 | SPP1 | IGHM | STX17-AS1 | SLCO5A1 | SLC11A1 | HDC |
| CD96 | NRP1 | ARHGAP24 | CSF2RA | NEGR1 | MXD1 | KIT |
| LEF1 | RGL1 | MS4A1 | GPAT3 | FLT3 | CD300E | PDE3A |
| ZBTB16 | SPRED1 | PLEKHG1 | SGK1 | HDAC9 | PLAUR | RIPOR3 |
| ETS1 | LGMN | FCRL1 | SLC8A1 | IRF8 | CSF3R | SLC24A3 |
| SAMD3 | AC131944.1 | KHDRBS2 | HLA-DPA1 | WNT5B | DYSF | AC068234.2 |
| MLLT3 | CD14 | OSBPL10 | HLA-DQB1 | PLEKHM3 | FGD4 | AC004083.1 |
| TXK | CD163 | PCDH9 | ENTPD1 | ZNF366 | KSR1 | PKHD1L1 |
| STAT4 | LYVE1 | MME | HLA-DQA1 | SHTN1 | MEGF9 | ITGA2B |
| IL7R | STAB1 | COL19A1 | AC104365.1 | ID2 | KYNU | GATA2 |
| NELL2 | SLC9A9 | TCF4 | CD74 | AC060234.3 | S100A9 | HPGD |
| ITK | COLEC12 | BCL11A | NR4A3 | SLC24A4 | AC084871.1 | RGS6 |
| FYN | ITSN1 | STRBP | SLC8A1-AS1 | IL15 | ARHGAP26 | ADCYAP1 |
| LINC01934 | NAV2 | KLHL14 | RTN1 | RAB7B | AQP9 | ANKRD55 |
| TC2N | SH3RF3 | GNG7 | STK38L | AC099560.1 | NLRP3 | ANXA1 |
| TOX | SLCO2B1 | LINC01374 | CCDC26 | VAC14 | LRRK2 | LINC00534 |
| TNIK | ABCA1 | PARP15 | SERPINB9 | IDO1 | GLUL | AGAP1 |
| SCML4 | FRMD4B | BLK | CTNND1 | ENOX1 | IL1B | CPA3 |
| RORA-AS1 | HRH1 | BLNK | CLEC7A | MIR924HG | TLR2 | AC006387.1 |
| TCF7 | DOCK4 | CD22 | PHC2 | ENPP1 | SAMSN1 | CNST |
| LTB | MS4A6A | ACSM3 | PLXDC2 | HLA-DPB1 | LYZ | PRKAR2B |
| IKZF2 | LRMDA | TPD52 | RGS1 | CLEC9A | S100A8 | SORBS1 |
| PRKCQ | ITPR2 | BACH2 | OGFRL1 | MCOLN2 | SLC2A3 | COL24A1 |
| BTBD11 | WDFY3 | GAB1 | VASH1 | SDK1 | IRAK3 | LMNA |
| CCND3 | SELENOP | DTX1 | CIITA | AC005599.1 | SLC43A2 | MEIS2 |
| ZNF831 | MERTK | ZCCHC7 | CST3 | VOPP1 | GLT1D1 | STON2 |
| CNOT6L | MS4A4A | RCSD1 | MGAM | CALCRL | IL1RAP | ARHGAP6 |
| LCK | FMN1 | AFF3 | GPR183 | CCDC26 | LYN | STXBP5 |
| PITPNC1 | ENPP2 | PRKCE | MALT1 | HLA-DQB1 | RBM47 | IL18R1 |
| GRAP2 | WWP1 | PLCG2 | GNA13 | CAMK2D | CD55 | CDH12 |
| PYHIN1 | TFRC | LRMP | INSIG1 | DAPP1 | ALDH1A2 | P2RX1 |
| ICOS | ADAP2 | RUBCNL | ITGAX | IDO2 | DPYD | ITGB3 |
| CD6 | STARD13 | IGKC | SULF2 | SLC22A23 | TNFRSF1B | ENPP3 |
| ANK3 | CCL3 | LINC00926 | AXL | KDM2B | FTH1 | EGF |
| PDE3B | ARHGAP18 | RHEX | PRKCA-AS1 | DNASE1L3 | ATP13A3 | PARD3 |
| SYTL2 | RBM47 | SP140 | IPCEF1 | FNIP2 | EMILIN2 | MAST4 |
| ADAMTS17 | MKNK1 | AL589693.1 | CLEC10A | CD74 | ADGRE2 | GLYATL1 |
| OXNAD1 | PDGFC | AP002075.1 | TYMP | NLGN4Y | DMXL2 | VCL |
| SATB1-AS1 | CCL4L2 | IGHD | RALA | NAV1 | FNDC3B | CXXC4-AS1 |
| SLFN12L | RNASE1 | CCDC50 | ADAM19 | AC007991.3 | ATP2B1 | AC073167.1 |
| INPP4B | NPL | SEL1L3 | PTGS2 | P2RY14 | FGR | LINC02197 |
| TOX2 | CTSB | ANGPTL1 | RUNX3 | FMNL2 | PTPRE | AL390957.1 |
| RETREG1 | DTNA | FAM214A | DOCK5 | ETV6 | FBXL5 | HPGDS |
| PTPN22 | PRDM1 | CLEC17A | AC087286.2 | KIF16B | GPCPD1 | PLPP1 |
| PTPN4 | EPB41L3 | PIP5K1B | CD83 | ANPEP | RTN1 | MITF |

| **Capillary Endothelial** | **Vasc Sm MM** | **Arterial Endothelial** | **Pericytes** |
| --- | --- | --- | --- |
| SELE | ACTA2 | DKK2 | GUCY1A2 |
| MCTP1 | ERBB4 | SERPINE2 | EBF2 |
| EMCN | MYH11 | MECOM | TEX41 |
| COL15A1 | CSMD1 | PCSK5 | EPS8 |
| ADGRL4 | TAGLN | SULF1 | ABCC9 |
| FLT1 | THSD4 | MGP | DLC1 |
| PCDH17 | SLIT3 | ARL15 | TRPC6 |
| APLNR | DMD | KCTD12 | LINC01091 |
| ARHGAP18 | CNNM2 | DEPP1 | PDGFRB |
| DACH1 | PRDM6 | SMAD6 | CCDC102B |
| RGCC | ALDH1A2 | EYS | ENPEP |
| C1orf112 | ANTXR1 | CXCL12 | NCKAP5 |
| FGD4 | MYOCD | SRGN | MYO1B |
| ADGRL2 | CACNA1C | PTPRB | EBF1 |
| FLI1 | CTNNA3 | GJA5 | PLCL1 |
| PRCP | NTRK3 | ITPR2 | SEMA5A |
| ANO2 | STAMBPL1 | ID1 | PRRX1 |
| KHDRBS2 | CCN2 | VEGFC | ZEB2 |
| PKP4 | KANK1 | ELN | SLC35F1 |
| PLVAP | LINC01099 | EXOC6 | ITGA1 |
| KIT | PDE3A | EFNB2 | GRID1 |
| PRKCH | GPC6 | CLEC14A | COL6A3 |
| NAV3 | PDLIM3 | HSPG2 | ARHGAP42 |
| NOSTRIN | PPP1R12B | ADAMTS6 | DPY19L2 |
| AC097459.1 | RCAN2 | ST8SIA6 | RERG |
| NEDD9 | ZFHX3 | BHLHE40 | EGFLAM |
| CD93 | SYNPO2 | EPAS1 | AC024901.1 |
| KITLG | PDLIM5 | SULT1E1 | LHFPL6 |
| RGS6 | SLC8A1 | TM4SF1 | GRK3 |
| ST8SIA4 | CDH13 | FBLN5 | GPM6B |
| VWF | FRY | ATP13A3 | DOCK10 |
| NRP1 | HMCN1 | SYT1 | KCNQ5 |
| PALMD | GNG2 | MMP16 | KHDRBS3 |
| CYYR1 | CADM2 | STOM | COL25A1 |
| PLEKHG1 | ADAMTSL1 | RNASE1 | EDNRA |
| NR5A2 | CARMN | PREX2 | NOTCH3 |
| RASAL2 | MYL9 | CALCRL | FAM162B |
| HPGD | SDK1 | GLUL | SEMA5B |
| DOCK4 | ZNF385B | BCAT1 | AC106845.1 |
| CEMIP2 | SLC26A7 | FBLN2 | PAG1 |
| ITGA6 | ELN | TMEM100 | AC008250.1 |
| KIAA1217 | UNC5C | PLPP1 | PDZD2 |
| ICAM1 | PXDNL | LMCD1 | TENM4 |
| EPAS1 | ST6GAL2 | CLDN5 | DCLK1 |
| SPC25 | PI15 | SRP14 | RAD51B |
| FENDRR | PDZRN3 | GFOD1 | SOX5 |
| RAPGEF5 | TPM1 | HEG1 | SLC24A3 |
| ETS2 | MYH10 | KLF2 | LINC02398 |
| PDE3B | ID4 | GJA4 | SLCO3A1 |
| AL356124.1 | LINC01098 | SMAD7 | AC022126.1 |

| **LGR5+ Basal 2** | **Secretory** | **LAP** | **Multiciliated 2** | **Bud Tip Adjacent** | **Bud Tip Progenitor** |
| --- | --- | --- | --- | --- | --- |
| TP63 | CP | ADAMTSL1 | DNAH12 | AC107223.1 | SFTPC |
| IGFBP2 | BPIFB1 | GPC5 | CFAP299 | GPC5 | DLG2 |
| KRT15 | RIMS1 | GDF15 | DNAH11 | AL132857.1 | ATP11A |
| LINC02057 | SCGB3A1 | AC005165.1 | NEK10 | ATP11A | PIK3C2G |
| MMP10 | SCGB3A2 | DAPK1 | TMEM232 | KHDRBS2 | ADAMTS6 |
| F3 | SLC4A4 | SCGB3A2 | CDHR3 | SFTPB | 1-Dec |
| ETS2 | SCGB1A1 | CLIC5 | CFAP54 | SFTA3 | RNF220 |
| ZSWIM6 | TLL1 | GLCCI1 | AGBL4 | CPM | HIF1A-AS3 |
| GPC3 | RARRES1 | CLIC6 | HYDIN | ROR1 | ROS1 |
| KRT17 | KDR | PLCL1 | DNAH9 | WIF1 | ETV5 |
| SPINK5 | FCGBP | DMD | STK33 | ADAMTSL1 | AC046195.2 |
| TIMP3 | CCL20 | SECISBP2L | DNAH7 | SMARCA5 | LINC01331 |
| NDFIP2 | IL1RAPL2 | CTNND2 | DCDC1 | AL356737.2 | NECTIN3 |
| MIR205HG | BMPR1B | MAGI3 | PACRG | LAMA3 | AC046195.1 |
| FN1 | LINC02300 | SEMA3E | CFAP47 | AC112206.2 | CD36 |
| IER3 | NAMPT | LMO3 | ZBBX | CTNNA3 | FAM184A |
| TSHZ2 | STEAP4 | CHN1 | SPAG17 | ST6GALNAC5 | ROR1 |
| PTPRZ1 | TMC5 | NEDD4L | LRRIQ1 | DLC1 | WIF1 |
| RAB30 | TFF3 | ERBB4 | DTHD1 | NDNF | SOX5 |
| IQCJ-SCHIP1 | CD55 | MBNL1 | PTPRT | SOX5 | ACOXL |
| PDLIM1 | AGR3 | ANOS1 | ERICH3 | MBIP | AL137009.1 |
| SLC6A6 | CFTR | GALNT13 | CFAP157 | GPC5-AS2 | C10orf90 |
| AC044810.2 | CA10 | TGFBR2 | DNAH6 | CD36 | MICAL2 |
| LMNA | LRMP | CFTR | DNAH3 | PHACTR1 | NHSL1 |
| NRXN3 | TMEM45A | CYB5A | C8orf34 | TMEFF2 | SFTA3 |
| LGR5 | NEGR1 | KHDRBS2 | DNAAF1 | P3H2 | LRMDA |
| AC026333.3 | LCN2 | PTPRD | ANKFN1 | ROS1 | HIF1A |
| DISC1 | HPGD | ZNF608 | CFAP43 | EMP2 | AL132857.1 |
| KRT14 | CXCL8 | DAPK2 | ADGB | AL139807.1 | CABLES1 |
| HMGA2 | ARHGEF38 | FAM189A2 | LMNTD1 | MECOM | TNC |
| AC084816.1 | CXCL2 | SFTPB | ARMC3 | LRRTM3 | FOXP2 |
| NCKAP5 | LYN | P3H2 | DNAH5 | ANKRD29 | PRICKLE1 |
| DST | CAPN8 | PLEKHG1 | ARMC4 | SLC22A3 | SEL1L3 |
| IL33 | ARFGEF3 | AL356737.2 | TTC29 | PTPRQ | NR3C2 |
| TENM4 | ERO1A | RALYL | VWA3A | LIMCH1 | PTPN13 |
| CDH3 | PIGR | UNC13C | AC130456.2 | AC073114.1 | TMEM163 |
| KLHL29 | NCOA7 | DSTN | SPAG16 | KCNJ15 | PKHD1 |
| LAMC2 | AHR | AC107223.1 | ULK4 | FILIP1 | HS2ST1 |
| MRTFB | SLC41A2 | CD47 | SPEF2 | CASC15 | AC073114.1 |
| LYPD6B | SFTPB | IL1RAPL2 | DNAH10 | CD47 | CELF2 |
| LINC00511 | TGM2 | HOPX | RSPH1 | ARHGAP44 | DLC1 |
| EGFR | LMO3 | MECOM | RNLS | AC046195.2 | MYO1B |
| KLHL13 | KYNU | AHR | FANK1 | FREM2 | SLC4A4 |
| KRT5 | PROM1 | WWC2 | KIAA2012 | FREM3 | DMBT1 |
| SLC20A2 | XBP1 | GPC5-AS2 | CFAP70 | AC046195.1 | TGFB2 |
| DPYSL3 | WSB1 | RBPMS | CEP126 | DAPK1 | THSD7A |
| GPRC5A | EPS8 | PRKCE | DNAI1 | PKHD1 | UGT2B7 |
| COL4A5 | PAPSS2 | CDKN1C | CFAP46 | MBNL1 | KCNJ15 |
| SEMA5A | AC003991.1 | KCNT2 | SPATA17 | PON2 | ABCA3 |
| SMOC2 | ADAMTSL1 | NCKAP5 | VWA3B | MAGI3 | NPC2 |

| **Epibasal** | **Serous** | **LGR5+ Basal 1** | **Multiciliated 1** | **Goblet** | **Proliferative basal** |
| --- | --- | --- | --- | --- | --- |
| EMP1 | SLC12A2 | COL1A2 | CFAP299 | BPIFB2 | MIR924HG |
| CSTB | EHF | COL1A1 | CDHR3 | MUC5B | DIAPH3 |
| KRT4 | CLDN10 | COL3A1 | CAPS | GOLM1 | SYNPO2 |
| H19 | NCALD | BNC2 | DNAH7 | AGR2 | TP63 |
| PLAT | CLDN10-AS1 | LDB2 | CFAP54 | DACH2 | BRIP1 |
| MUC4 | KCNMA1 | NFATC2 | STK33 | CNTD1 | KRT17 |
| AQP3 | SLCO1A2 | COL11A1 | LRRIQ1 | ST6GALNAC3 | KRT14 |
| DENND2C | ITPR2 | COL14A1 | DNAH12 | AF233439.1 | TIMP3 |
| KRT6A | CCL28 | ZFPM2 | PIEZO2 | UAP1 | MIR205HG |
| SERPINB13 | CHRM3 | COL5A2 | FAM155A | NKX3-1 | COL14A1 |
| CLDN4 | PIP5K1B | ZNF385D | AGBL4 | B4GALT1 | FRMD4B |
| KRT13 | BARX2 | COL2A1 | DNAH11 | FGF13 | CALD1 |
| SERPINB5 | FCGBP | DCN | PTPRT | ATP2A3 | CENPF |
| CDH26 | ESRRG | POSTN | NEK10 | CMTM8 | COL4A2 |
| TACSTD2 | PDE4B | COL9A1 | TMEM190 | ACSL3 | FGFR1 |
| MIR4435-2HG | ITPRID1 | FLRT2 | CFAP157 | GMDS | CENPP |
| ALDH1A3 | STK39 | PLXDC2 | DNAH9 | GFPT1 | EGFR |
| AC015712.2 | LTF | PRKG1 | AC130456.2 | PTPRN2 | TOP2A |
| IGFBP3 | LHFPL2 | LSAMP | SPAG17 | INHBA | PTPN14 |
| ANXA1 | TNFRSF19 | EBF1 | TMEM232 | NR4A2 | AC044810.2 |
| CLDN1 | PKN2-AS1 | SPARC | HYDIN | MLPH | NFIB |
| APOBEC3A | CARMIL1 | TENM3 | PACRG | ODC1 | MEG8 |
| PLAUR | C4orf19 | SGCD | DNAH3 | BAIAP2 | MEG3 |
| CERNA2 | TM4SF1 | COL12A1 | ZBBX | ANKRD37 | VCAN |
| KRT5 | GMDS | GPC6 | DNAAF1 | MSMB | ATAD2 |
| KRT19 | CRACR2A | CCDC102B | DNAH6 | DGKD | MYO3A |
| PLAC8 | SLC24A3 | NAV3 | RSPH1 | MIA3 | KIF23 |
| S100A2 | GNAS | ZFPM2-AS1 | C8orf34 | COBL | ANKH |
| LINC00511 | ALDH1A3 | ACAN | DCDC1 | BACE2 | NCKAP5 |
| SERPINB2 | AL139383.1 | ZEB1 | ERICH3 | TNFRSF19 | BOC |
| ITPKC | NFKBIZ | COL6A6 | DNAH5 | XBP1 | SYN3 |
| ADGRF1 | DEPTOR | ADAMTSL3 | ARMC4 | TPM4 | SEMA5A |
| NEAT1 | INHBA | COL5A1 | CFAP47 | SLIT3 | RNF19A |
| EPHA2 | DTNB | PIEZO2 | ZNF385D | GALNT7 | BCL2 |
| ANXA2 | KANK1 | FBN1 | CFAP43 | FAM83D | PLCB1 |
| DSP | KIAA1324 | HPSE2 | DTHD1 | CHRM3 | ROBO1 |
| RAB11FIP1 | ERBB4 | CDH11 | VWA3A | ERN2 | ECT2 |
| SERPINB1 | SLC5A1 | FBLN1 | LMNTD1 | ADARB2 | LRP4 |
| ST14 | ADGRV1 | IL1RAPL1 | AL139815.1 | CREB3L1 | ASAP1 |
| ATP12A | INSR | GALNT17 | ADGB | PLCB4 | PDZD2 |
| LYN | PALMD | PRRX1 | CFAP73 | ACLY | GHR |
| RAP2B | MAP2 | COL24A1 | GNA14-AS1 | OPCML | VCL |
| TNNT3 | SLPI | MIR100HG | CEP126 | ANXA1 | EYA2 |
| IL1RN | COL28A1 | ABI3BP | ULK4 | KIAA1324 | IQCJ-SCHIP1 |
| UBC | XBP1 | MEG8 | RNLS | GALNT12 | RFC3 |
| CYP2C18 | PLCB1 | PLCB1 | SPEF2 | DOP1B | RARB |
| ID1 | PACS1 | EPB41L2 | ERICH3-AS1 | ACVR1 | RBBP8 |
| S100A6 | SORBS2 | FAT4 | ARMC3 | CBR3 | MPPED2 |
| PKP1 | ITGA6 | MEG3 | DNAH10 | AL139383.1 | KIAA1217 |
| MACC1 | ELMO1 | LAMA4 | ANKFN1 | ARHGAP26 | RRM2 |

| **Transitional basal** | **GRP+ PNEC** | **GHLR+ PNEC** | **Deuterosome** | **GAP43+ PNEC** |
| --- | --- | --- | --- | --- |
| CERNA2 | GRP | GHRL | CDC20B | KCNIP4 |
| KRT5 | AC109466.1 | ACSL1 | CCNO | CNTNAP2 |
| KRT6A | NRG1 | NRXN1 | CFAP47 | DSCAM |
| EMP1 | NKAIN2 | RFX6 | GPX8 | CNTN5 |
| KRT15 | RIMS2 | RIMS2 | AGBL4 | LSAMP |
| IGFBP3 | RALYL | CPE | CCDC171 | LRP1B |
| LINC02057 | CACNA1A | RGS7 | CFAP299 | CTNNA2 |
| LINC00511 | RGS7 | PEX5L | DNAH12 | SYT1 |
| MIR205HG | NRXN1 | CACNB2 | CEP112 | ALK |
| PLAT | ATP8A2 | RIMBP2 | LRRIQ1 | KCNQ5 |
| S100A2 | AL589693.1 | AC084871.1 | CROCC2 | UNC5C |
| ADAM28 | RBFOX1 | GRIK1 | CEP128 | RBFOX1 |
| EDN2 | LINGO2 | KCNMB2-AS1 | PLOD2 | ANK2 |
| SDK1 | SYT1 | NPAS3 | DEUP1 | FGF13 |
| ETS2 | KCNB2 | DPP10 | ADGB | HS6ST3 |
| PDLIM1 | CPE | TOX | LINC01091 | DLGAP2 |
| AC026333.3 | NOL4 | RAP1GAP2 | HYDIN | DPP6 |
| LAMC2 | CACNA2D3 | ANK2 | SPAG16 | EML5 |
| TACSTD2 | CADM2 | SGCZ | PPM1E | RYR2 |
| ZSWIM6 | ST18 | PDE1C | DNAH7 | PLXNA4 |
| PTPRZ1 | AC008415.1 | TMEM108 | STRBP | ZNF804A |
| AC084816.1 | MIAT | CACNA2D1 | WDR49 | AC092691.1 |
| SLC25A25 | ANK2 | AL589693.1 | SPAG17 | SEMA6D |
| H19 | FAM155A | EYA4 | ARMC3 | LINGO2 |
| IER3 | CADPS | ABCC8 | PRMT8 | MGAT4C |
| UBC | DGKB | SMOC1 | STIL | SYN2 |
| NT5DC3 | TMEM178B | GPC6 | CFAP54 | NCAM2 |
| F3 | SPHKAP | KCNB2 | PLK4 | SPOCK3 |
| TP63 | SEC11C | KCNMB2 | TMEM232 | HDAC9 |
| PCDH7 | SCN3A | NPSR1-AS1 | KIF24 | AC009264.1 |
| ID1 | GRIK2 | CFC1 | SPATA17 | AC110296.1 |
| AQP3 | KCNMB2-AS1 | HEPACAM2 | HSP90AA1 | PRUNE2 |
| KRT17 | SRRM4 | RAPGEF4 | NEK10 | CHRM2 |
| AC120193.1 | GALNT13 | PDE8B | E2F7 | SLC8A1 |
| GLIS3 | MGAT4C | PROX1 | EZH2 | AC093879.1 |
| CD9 | AC019197.1 | GRIK2 | DCDC1 | ENOX1 |
| AC104123.1 | DTNA | C12orf75 | MYB | SGCZ |
| EZR | IGFBP5 | GABRB3 | CEP83 | PLD5 |
| GPRC5A | DPP10 | PPP2R2B | FGFR1OP | NCAM1 |
| LINC00621 | NPAS3 | DSCAML1 | AL139815.1 | ADGRB3 |
| KRT19 | PIEZO2 | FSTL5 | FOXN4 | STMN2 |
| LYPD6B | STXBP5L | COL3A1 | ULK4 | MAP1B |
| JUNB | ROBO1 | FAM155A | DNAH6 | ATRNL1 |
| PKP1 | SLC8A1 | SEMA3A | AC234582.1 | NELL2 |
| EGFR | KCNMB2 | PCSK1N | LINC00970 | PCDH15 |
| TNFAIP8 | UNC5D | SLC8A1 | PCM1 | MIR137HG |
| NTN1 | CNTNAP2 | AFF3 | C3orf67 | MYT1L |
| SLC2A1 | PTPRN2 | UNC80 | NPHP1 | DGKI |
| KRT13 | GRIA2 | SLC4A10 | RGS22 | PPP2R2B |
| ARHGAP24 | CDH18 | SORL1 | FOXJ1 | BNC2 |

| **RSPO2+ Mes 1** | **RSPO2+ Mes 2** | **Twist2+ Prox Mes** | **Chondroblast** | **RSPO2+ Mes 3** | **SCARA5+ Mes** |
| --- | --- | --- | --- | --- | --- |
| FAM155A | NKAIN2 | CSMD1 | COL1A1 | CNTNAP2 | GPC5 |
| MACF1 | UNC13C | ERBB4 | KCNIP4 | AL136456.1 | ADAMTSL1 |
| ADH1B | FAT3 | MACROD2 | NTRK2 | NKAIN2 | NCAM2 |
| PLEKHH2 | ARHGAP28 | LTBP1 | COL1A2 | GLIS3 | COL3A1 |
| PCDH15 | NTM | RGS6 | SPARC | CCBE1 | SEMA3C |
| TCF21 | TENM2 | ELN | COL3A1 | ALDH1A2 | BRINP3 |
| GABRG3-AS1 | RSPO2 | ADAMTS19 | ARHGAP26 | SLIT2 | SLC26A7 |
| PRKCB | SLC4A4 | TWIST2 | DCLK1 | FAM155A | CNTN4 |
| A2M | FREM1 | AC008591.1 | GAS1 | PRKCB | PID1 |
| CCBE1 | ADGRB3 | MECOM | MIR99AHG | TMEM108 | ADAMTSL3 |
| DACH1 | GABRG3-AS1 | MME | PTN | FREM1 | GRID2 |
| CTTNBP2 | GABRG3 | PHACTR1 | ASPN | ADH1B | AKAP12 |
| ELMO1 | SLFN12L | SLC6A2 | SULF1 | BX284613.2 | PI15 |
| GABRG3 | PRKCB | TMTC1 | CDON | TCF21 | COL1A1 |
| LURAP1L-AS1 | CCBE1 | FGF13 | SVIL | CNTN1 | GPC6 |
| NPNT | ENAH | SLC8A1 | EPHA7 | SELENOP | SPARC |
| GALNT13 | PDE3B | NBEA | EBF2 | MROH9 | FLRT2 |
| ITGA8 | ROBO2 | NAV3 | SFRP2 | SEMA3D | TENT5A |
| SLIT2 | THSD7A | DOK5 | FOXP1 | CDH12 | MFAP5 |
| NTNG1 | MYO10 | COL6A6 | GFRA1 | PIEZO2 | LINC01197 |
| MYLK | KALRN | ZEB1 | GAS2 | C7 | ADAMTS19 |
| LMO4 | AC025508.1 | ATXN1 | BCKDHB | DAPK1 | ZFPM2 |
| CNTNAP2 | TMEM178B | VPS13B | TENT5A | DACH1 | LINC00924 |
| RSPO2 | ITGA8 | LRRC7 | IGF1 | A2M | CACNB4 |
| DOCK4 | DOCK4 | POSTN | LGR4 | BTG1 | AL354771.1 |
| FAT3 | GUCY1A1 | MIR99AHG | CCDC80 | KCNMB2 | EBF1 |
| SELENOP | STXBP6 | AHR | LRRC7 | MEOX2 | SULT1E1 |
| ATP8A1 | LYPD6B | PAPPA | COL6A6 | FRAS1 | NAV3 |
| NR2F1 | DACH1 | VSTM4 | MGP | CNTN3 | CCDC102B |
| AC005550.2 | ANK3 | CREB5 | COL5A1 | GPC3 | COL1A2 |
| BTG1 | HAS2 | DMD | OGN | CNKSR2 | AC008591.1 |
| CDH12 | BMPER | RBPJ | KLF4 | MYLK | SEMA3D |
| LIMD1 | PLEKHH2 | COL4A6 | EBF3 | THBS1 | SCARA5 |
| PPP1R3C | GRK3 | TIMP3 | COL12A1 | PCDH7 | FBLN1 |
| CDH4 | SNCA | OSR2 | NTRK3 | ZDHHC14 | NTRK3 |
| SLC22A23 | TMEM132C | PDE7B | PRSS23 | 1-Mar | DCHS2 |
| LURAP1L | GALNT13 | COL4A5 | EIF4A3 | RSPO2 | SERPINF1 |
| CNTN3 | AFF3 | SULF1 | ALCAM | PREX2 | STAC |
| FRAS1 | BMP5 | ZNF536 | KLF2 | MYO10 | ARHGEF3 |
| C7 | KIF26B | PGM5 | SERPINF1 | DCC | MID1 |
| MYO10 | FAM155A | AL355612.1 | MACROD2 | VEGFC | MSC-AS1 |
| PTPRD | PTCH1 | HPSE2 | DLK1 | LINC02224 | FBN1 |
| DST | ROR1-AS1 | FAT4 | PLAC9 | NR2F1 | ZFPM2-AS1 |
| FILIP1 | TBX2 | AC124852.1 | PI15 | SAT1 | CERT1 |
| AFDN | AC124854.1 | TLE4 | DOCK11 | CACNB2 | LINC02643 |
| DCC | PIEZO2 | SGIP1 | PRRX1 | KAZN | TGFBR2 |
| SCN7A | HECTD2 | LDB2 | THBS2 | AFDN | IGFBP5 |
| KALRN | PDE4D | CNKSR3 | SLC41A2 | ADAMTS12 | DPYSL3 |
| AC025508.1 | C7 | PLCB4 | DCN | RYR2 | TRIO |
| ANGPT1 | GNAI1 | LINC00578 | DPYSL3 | KCNMB2-AS1 | KLF4 |

| **Fibroblast** | **RSPO2+ Mes 4** | **Ptch2+ Mes 2** | **Prolif. Mes. 1** | **Chondrycyte** | **Myofibroblast** |
| --- | --- | --- | --- | --- | --- |
| ADAM12 | FAM155A | KCND2 | DIAPH3 | COL2A1 | DACH2 |
| NAV3 | TCF21 | IGFBP7 | BRIP1 | COL9A1 | SLC4A4 |
| CCDC102B | ADH1B | SEMA5A | EZH2 | ACAN | SYT1 |
| EBF1 | SELENOP | MSC-AS1 | MIR924HG | WWP2 | PLPPR4 |
| AHR | RSPO2 | GEM | CENPK | COL11A1 | ATRNL1 |
| ADAMTS19 | GALNT13 | MGAT4C | ATAD2 | PLCB1 | PRAG1 |
| SEMA5A | PRKCB | CNTN4 | TOP2A | SOX6 | MYOCD |
| ABCA10 | NKAIN2 | LINC01197 | APOLD1 | SORBS2 | CCDC68 |
| CSMD1 | FAT3 | LINC00924 | RRM2 | COL11A2 | SOX5 |
| AC008591.1 | DACH1 | NPR3 | MELK | COL9A2 | SEMA3A |
| DCLK1 | A2M | PDLIM3 | CENPP | HAPLN1 | TANC1 |
| SULT1E1 | FRAS1 | COL4A1 | NUSAP1 | SOX5 | PTCH1 |
| GPC5 | CDH12 | PTCH2 | HELLS | LRP1B | PAG1 |
| EBF2 | SLIT2 | FRMPD4 | C21orf58 | NFATC2 | MCTP2 |
| TENM4 | ARHGAP28 | CCDC102B | SMC4 | COL27A1 | CPED1 |
| POSTN | MYLK | GPC6 | LINC01572 | STK32B | ADGRB3 |
| LAMB1 | CCBE1 | LINC00578 | KNL1 | COL9A3 | LMCD1 |
| FGF13 | CNKSR2 | FHOD3 | FANCI | PRDM16 | ANO4 |
| KLHL29 | THSD7A | SPARC | ASPM | CHST11 | ACTA2 |
| TENM1 | GABRG3 | JAG1 | RFC3 | NR4A2 | EYA4 |
| MID1 | AL136456.1 | CGNL1 | SPC25 | FRY | SNED1 |
| KLHL13 | CNTNAP2 | COL3A1 | POLQ | XYLT1 | LEF1 |
| LDB2 | FREM1 | NIBAN1 | KIF15 | HDAC9 | ANO3 |
| EBF3 | MEOX2 | KCNIP1 | NSD2 | SORBS1 | SDC2 |
| MAGI1 | ITGA8 | BX322234.1 | CENPF | BMPR1B | FENDRR |
| EDNRA | UNC13C | ASPN | DTL | ABTB2 | ZNF536 |
| AL355612.1 | TMSB10 | RTN1 | MKI67 | SIM2 | ROBO2 |
| CASC15 | PLEKHH2 | NKAIN3 | CIT | NR4A1 | AC005358.1 |
| TBX18 | LYPD6B | GRID2 | KIF11 | FGFR2 | MAML3 |
| TWIST2 | MACF1 | THBS2 | KNTC1 | SDK2 | COL24A1 |
| DKK2 | C7 | ZFPM2 | BARD1 | TAFA2 | ARHGAP6 |
| CFH | DOCK4 | LRRC7 | NCAPG2 | KCNQ5 | KIF26B |
| LHFPL2 | PIEZO2 | DLGAP1 | ZGRF1 | ADCY2 | CARMN |
| CLCN5 | WNT2 | STAC | TPX2 | TOX | TMEM132C |
| ABCA9 | NTNG1 | CASC15 | FANCA | SCGB3A1 | ANKS1B |
| LHFPL6 | FN1 | SPON2 | ECT2 | ERG | HIPK2 |
| TLN2 | NPNT | LHFPL6 | MMS22L | PCAT1 | CHN1 |
| HPSE2 | ANGPT1 | DMD | TMPO | MEG3 | NKAIN2 |
| PDGFRB | GLIS3 | EPHA3 | PRIM2 | APBB2 | MYH11 |
| SNTB1 | LURAP1L | KANK1 | ATAD5 | MYO1D | UNC13C |
| PAWR | BTG1 | NPAS2 | CENPU | EDIL3 | ST6GALNAC3 |
| ITM2A | TMEM108 | AC079142.1 | ANLN | ADGRA3 | SLFN12L |
| PTCH2 | TFPI | ZFPM2-AS1 | NCAPD3 | CSGALNACT1 | SEMA3C |
| COL6A6 | MYO10 | ARHGAP15 | HMGB2 | PEG3 | NKD1 |
| ANK2 | AFDN | MMP16 | DLEU2 | FAM160A1 | AC005237.1 |
| ABCA9-AS1 | GUCY1A1 | COL4A2 | BRCA1 | TFRC | TBX5 |
| PHACTR1 | ANK3 | AC064875.1 | LMNB1 | GRIK2 | PDGFRA |
| BX284613.2 | ELMO1 | DPYSL3 | MIS18BP1 | TENM3 | SNHG14 |
| ST8SIA1 | MDK | MCC | NCAPG | PDGFC | IRS1 |
| TCF7L2 | CNTN3 | ADAM12 | TUBA1B | PTPRZ1 | ETV5 |

| **Chondrocyte Precursor** | **Mes. Cluster 13** | **Ptch2+ Mes 1** | **Mature Chondrocyte** | **Smooth Muscle** | **Prolif. Mes 2** |
| --- | --- | --- | --- | --- | --- |
| COL8A1 | COL25A1 | EBF2 | COL9A1 | HHIP | ASPM |
| TENM3 | MECOM | AC116345.1 | COL11A1 | MYH11 | TOP2A |
| MPPED2 | TENM3 | PTCH2 | COL2A1 | AC098588.3 | CENPF |
| CRISPLD1 | PTN | SLC22A3 | WWP2 | ACTA2 | DIAPH3 |
| COL12A1 | PDE3A | MATN2 | ACAN | SYNPO2 | CENPE |
| KHDRBS2 | EYA4 | AL590807.1 | XYLT1 | DOCK3 | MKI67 |
| FGFR2 | KHDRBS2 | RGS6 | SORBS2 | CARMN | ARL6IP1 |
| COL24A1 | MPPED2 | SNTG2 | ZNF385B | MYOCD | APOLD1 |
| LMO7 | NTRK2 | AHR | COL11A2 | ACTG2 | HMGB2 |
| CHL1 | MACROD2 | LINGO2 | SOX6 | NTN1 | SMC4 |
| ADAMTS17 | TRPS1 | NTRK3 | COL9A2 | ZNF536 | TPX2 |
| TOX | SPOCK3 | DCX | COL27A1 | SEMA3C | GAS2L3 |
| ARHGEF28 | FOXP1 | ABCA9 | EPYC | GREM2 | NAV2 |
| TRPS1 | THBS2 | ABCA10 | FRY | RBPMS | NUSAP1 |
| FARP1 | COL12A1 | TENM2 | HAPLN1 | AC005358.1 | ECT2 |
| AL122014.1 | SVIL | DKK2 | LRP1B | KCNMA1 | CCNB1 |
| NR4A1 | COL14A1 | BNC2 | SORBS1 | AC025280.3 | KNL1 |
| COL14A1 | PLCB4 | PCBP3 | CHST11 | TACC2 | CKAP2 |
| EYA4 | AC138627.1 | ADAM12 | CSGALNACT1 | CRISPLD2 | PRC1 |
| ZNF521 | PRR16 | MGAT5 | COL9A3 | ATRNL1 | KPNA2 |
| PDE3A | PRRX1 | PAK3 | SOX5 | TAGLN | TUBA1C |
| STK32B | GAS2 | SLC2A3 | SIM2 | DMD | ANLN |
| OPHN1 | FOXP2 | TLL1 | PLCB1 | HPSE2 | KIF23 |
| MECOM | KCNIP4 | NEGR1 | BMPR1B | ADAMTS1 | CDK1 |
| FOXP1 | SLC8A1 | KCNIP1 | PRDM16 | ADAMTS6 | KIF11 |
| HDAC9 | PDGFD | EGFR | TAFA2 | TPM1 | TUBA1B |
| CTNNA3 | LMO7 | TENM4 | MEG3 | COL4A5 | SGO2 |
| BICC1 | KCTD1 | SYT1 | CNMD | LUZP2 | TUBB4B |
| FLI1 | LAMA3 | VIT | NR4A2 | PLXNA4 | CALM2 |
| THBS2 | EPHA7 | SPOCK1 | NFATC2 | DACH2 | CKAP5 |
| THSD4 | WNT5B | TGFBI | PCAT1 | ARHGAP42 | DLGAP5 |
| RAI14 | IQGAP2 | GAB2 | MATN1 | ENTPD1-AS1 | AC073529.1 |
| NPAS3 | PPP3CA | MAGI1 | AL022068.1 | MRVI1 | KIF14 |
| PDZD2 | DOCK11 | ADAMTS14 | SDK2 | BRINP3 | NUF2 |
| TIAM2 | ITGBL1 | ANK2 | GRIK2 | IGFBP5 | BUB1B |
| ERG | BX322234.1 | PELI2 | PEG3 | DPP10 | ARHGAP11B |
| ITGBL1 | ZBTB7C | SNTB1 | ITGA6 | COL4A6 | KIF18A |
| SPOCK3 | OPHN1 | ARHGAP42 | ENPP1 | LINC01322 | H2AFZ |
| PCLO | NR4A1 | EBF3 | MSR1 | STAMBPL1 | BUB1 |
| PRRX1 | EPHA6 | PAWR | AC011586.2 | SORBS1 | KIF4A |
| ANOS1 | NEXMIF | LINC01482 | ITIH6 | ADAM19 | GTSE1 |
| SMOC2 | PRSS23 | CNTN4 | PTPRZ1 | ZNF385D | KIF20B |
| GNAS | ZEB1 | JAG1 | C11orf58 | LPP | LINC01572 |
| FNDC1 | ADGRA3 | LEPR | FAM160A1 | PDGFC | NDC80 |
| MACROD2 | BICC1 | TBX18 | PDZD2 | LMOD1 | G2E3 |
| KCNQ5 | ARHGAP12 | PI16 | UCMA | AL354771.1 | PTTG1 |
| ABTB2 | EBF3 | RIPOR2 | NR4A1 | LRIG1 | LGALS1 |
| XKR6 | LDLRAD4 | YBX3 | LINC00511 | ZNF385D-AS2 | UBE2S |
| NEXMIF | PDZD2 | ABCA9-AS1 | EDIL3 | TBX3 | CKS2 |
| ADCY2 | LTBP1 | EGR3 | STK32B | FBXO32 | KIF18B |
