## Supplemental Table 2 for "A spatially-resolved blueprint of the developing human lung reveals a WNT-driven niche for basal stem cells"

| Gene | Ensembl ID | Probesets |
| --- | --- | --- |
| CDH1 | ENSG00000039068 | 8 |
| EPCAM | ENSG00000119888 | 8 |
| NKX2-1 | ENSG00000136352 | 8 |
| SOX2 | ENSG00000181449 | 8 |
| SOX9 | ENSG00000125398 | 8 |
| AGER | ENSG00000204305 | 8 |
| RTKN2 | ENSG00000182010 | 8 |
| MYRF | ENSG00000124920 | 8 |
| NTM | ENSG00000182667 | 5 |
| AQP5 | ENSG00000161798 | 8 |
| SFTPA1 | ENSG00000122852 | 5 |
| ZNF385B | ENSG00000144331 | 8 |
| LAMP3 | ENSG00000078081 | 8 |
| FMO5 | ENSG00000131781 | 8 |
| SLCO4C1 | ENSG00000173930 | 8 |
| SLC6A14 | ENSG00000268104 | 8 |
| RND1 | ENSG00000172602 | 8 |
| LRP2 | ENSG00000081479 | 8 |
| KRT5 | ENSG00000186081 | 8 |
| KRT17 | ENSG00000128422 | 8 |
| TP63 | ENSG00000073282 | 8 |
| SLC5A7 | ENSG00000115665 | 8 |
| KISS1 | ENSG00000170498 | 3 |
| MMP10 | ENSG00000166670 | 8 |
| CXCL1 | ENSG00000163739 | 8 |
| GABRB2 | ENSG00000145864 | 8 |
| GABRA1 | ENSG00000022355 | 8 |
| CDC20B | ENSG00000164287 | 8 |
| FOXJ1 | ENSG00000129654 | 8 |
| ZBBX | ENSG00000169064 | 8 |
| RSPH1 | ENSG00000160188 | 8 |
| CFAP73 | ENSG00000186710 | 8 |
| C6 | ENSG00000039537 | 8 |
| MUC16 | ENSG00000181143 | 8 |
| SPDEF | ENSG00000124664 | 8 |
| MUC5AC | ENSG00000215182 | 8 |
| MUC5B | ENSG00000117983 | 8 |
| NKX3-1 | ENSG00000167034 | 8 |
| LTF | ENSG00000012223 | 8 |
| BARX2 | ENSG00000043039 | 8 |
| CLDN10-AS1 | ENSG00000223392 | 6 |
| CCL28 | ENSG00000151882 | 8 |
| MUC4 | ENSG00000145113 | 8 |
| GDF15 | ENSG00000130513 | 8 |
| RNASE1 | ENSG00000129538 | 8 |
| GRP | ENSG00000134443 | 6 |
| ASCL1 | ENSG00000139352 | 8 |
| GHRL | ENSG00000157017 | 8 |
| CHGA | ENSG00000100604 | 8 |
| CALCA | ENSG00000110680 | 8 |
| PROX1 | ENSG00000117707 | 8 |
| NRP2 | ENSG00000118257 | 8 |
| RELN | ENSG00000189056 | 8 |
| STAB2 | ENSG00000136011 | 8 |
| DKK2 | ENSG00000155011 | 8 |
| FGF18 | ENSG00000156427 | 8 |
| BMX | ENSG00000102010 | 8 |
| IGFBP3 | ENSG00000146674 | 8 |
| FCN3 | ENSG00000142748 | 7 |
| SYN2 | ENSG00000157152 | 8 |
| KIT | ENSG00000157404 | 8 |
| SPC25 | ENSG00000152253 | 8 |
| APLNR | ENSG00000134817 | 8 |
| ACKR1 | ENSG00000213088 | 8 |
| VCAM1 | ENSG00000162692 | 8 |
| PLA1A | ENSG00000144837 | 8 |
| NOS1 | ENSG00000089250 | 8 |
| ANO2 | ENSG00000047617 | 8 |
| PECAM1 | ENSG00000261371 | 8 |
| VWF | ENSG00000110799 | 8 |
| HPGD | ENSG00000164120 | 8 |
| EDNRB | ENSG00000136160 | 8 |
| SOSTDC1 | ENSG00000171243 | 8 |
| CDH13 | ENSG00000140945 | 8 |
| APLN | ENSG00000171388 | 8 |
| LAMC3 | ENSG00000050555 | 8 |
| PDGFRB | ENSG00000113721 | 8 |
| FHL5 | ENSG00000112214 | 8 |
| CDH19 | ENSG00000071991 | 8 |
| MYOCD | ENSG00000141052 | 8 |
| LUZP2 | ENSG00000187398 | 8 |
| ZNF536 | ENSG00000198597 | 8 |
| FGF14 | ENSG00000102466 | 8 |
| SFRP2 | ENSG00000145423 | 8 |
| MFAP5 | ENSG00000197614 | 8 |
| PTCH2 | ENSG00000117425 | 8 |
| PI15 | ENSG00000137558 | 8 |
| GFRA1 | ENSG00000151892 | 8 |
| MOXD1 | ENSG00000079931 | 8 |
| SYN3 | ENSG00000185666 | 8 |
| RSPO2 | ENSG00000147655 | 8 |
| RANBP3L | ENSG00000164188 | 8 |
| UBA52 | ENSG00000221983 | 8 |
| OLFML3 | ENSG00000116774 | 8 |
| PSAP | ENSG00000197746 | 8 |
| RUNX1 | ENSG00000159216 | 8 |
| 1-Dec | ENSG00000173077 | 5 |
| CCL2 | ENSG00000108691 | 8 |
| PTPRC | ENSG00000081237 | 8 |
| KLRF1 | ENSG00000150045 | 8 |
| NCR1 | ENSG00000189430 | 8 |
| PRF1 | ENSG00000180644 | 8 |
| SH2D1B | ENSG00000198574 | 8 |
| MS4A1 | ENSG00000156738 | 8 |
| BANK1 | ENSG00000153064 | 8 |
| IGHM | ENSG00000211899 | 4 |
| PAX5 | ENSG00000196092 | 8 |
| SLC18A2 | ENSG00000165646 | 8 |
| MS4A2 | ENSG00000149534 | 8 |
| FCN1 | ENSG00000085265 | 8 |
| APOBEC3A | ENSG00000128383 | 8 |
| ADGRE2 | ENSG00000127507 | 8 |
| CD300E | ENSG00000186407 | 8 |
| CLEC4C | ENSG00000198178 | 8 |
| KCNK10 | ENSG00000100433 | 8 |
| CUX2 | ENSG00000111249 | 8 |
| CD6 | ENSG00000013725 | 8 |
| THEMIS | ENSG00000172673 | 8 |
| ITK | ENSG00000113263 | 8 |
| LEF1 | ENSG00000138795 | 8 |
| CD28 | ENSG00000178562 | 8 |
| CD4 | ENSG00000010610 | 8 |
| CD8A | ENSG00000153563 | 8 |
| APOE | ENSG00000130203 | 5 |
| CTSA | ENSG00000064601 | 8 |
| ATP6V0C | ENSG00000185883 | 8 |
| C1QC | ENSG00000159189 | 8 |
| MARCO | ENSG00000019169 | 8 |
| SIGLEC1 | ENSG00000088827 | 8 |
| CD163L1 | ENSG00000177675 | 8 |
| F13A1 | ENSG00000124491 | 8 |
| LILRB5 | ENSG00000105609 | 8 |
| STAB1 | ENSG00000010327 | 8 |
| FAM9B | ENSG00000177138 | 8 |
| GLDN | ENSG00000186417 | 8 |
| CLEC10A | ENSG00000132514 | 8 |
| CD1C | ENSG00000158481 | 8 |
| ZNF366 | ENSG00000178175 | 8 |
| CLEC9A | ENSG00000197992 | 8 |
| TUBB3 | ENSG00000258947 | 8 |
| CALB1 | ENSG00000104327 | 8 |
| SLC17A6 | ENSG00000091664 | 8 |
| TAC1 | ENSG00000006128 | 8 |
| TRPV1 | ENSG00000196689 | 8 |
| ADCYAP1 | ENSG00000141433 | 8 |
| GAP43 | ENSG00000172020 | 8 |
| DSCAM | ENSG00000171587 | 8 |
| CHAT | ENSG00000070748 | 8 |
| CD38 | ENSG00000004468 | 8 |
| SDC1 | ENSG00000115884 | 8 |
| IGHG1 | ENSG00000211896 | 7 |
| ACTA2 | ENSG00000107796 | 6 |
| MYH11 | ENSG00000133392 | 8 |
| FST | ENSG00000134363 | 8 |
| PLIN2 | ENSG00000147872 | 8 |
| ACTG2 | ENSG00000163017 | 8 |
| CNN1 | ENSG00000130176 | 8 |
| ASPN | ENSG00000106819 | 8 |
| CD34 | ENSG00000174059 | 8 |
| CSPG4 | ENSG00000173546 | 8 |
| SFTPD | ENSG00000133661 | 8 |
| GREM2 | ENSG00000180875 | 8 |
| DOCK3 | ENSG00000088538 | 8 |
| CXCL14 | ENSG00000145824 | 8 |
| ANO3 | ENSG00000134343 | 8 |
| KCNK17 | ENSG00000124780 | 8 |
| LDB3 | ENSG00000122367 | 8 |
| JPH2 | ENSG00000149596 | 8 |
| IL7R | ENSG00000168685 | 8 |
| PLA2G5 | ENSG00000127472 | 8 |
| PIEZO2 | ENSG00000154864 | 8 |
| COL11A1 | ENSG00000060718 | 6 |
| VIT | ENSG00000205221 | 8 |
| SPOCK1 | ENSG00000152377 | 8 |
| SHOX2 | ENSG00000168779 | 8 |
| KCND2 | ENSG00000184408 | 8 |
| NPR3 | ENSG00000113389 | 8 |
| GEM | ENSG00000164949 | 8 |
| SMOC1 | ENSG00000198732 | 8 |
| ITGA11 | ENSG00000137809 | 8 |
| COL8A1 | ENSG00000144810 | 8 |
| EPYC | ENSG00000083782 | 8 |
| CNMD | ENSG00000136110 | 8 |
| SLC6A2 | ENSG00000103546 | 8 |
| OSR2 | ENSG00000164920 | 8 |
| TWIST2 | ENSG00000233608 | 8 |
| MME | ENSG00000196549 | 8 |
| IL1RAPL2 | ENSG00000189108 | 8 |
| CP | ENSG00000047457 | 8 |
| SCGB3A2 | ENSG00000164265 | 7 |
| BPIFB1 | ENSG00000125999 | 8 |
| CAPN13 | ENSG00000162949 | 8 |
| CLIC5 | ENSG00000112782 | 8 |
| SLC34A2 | ENSG00000157765 | 8 |
| DMBT1 | ENSG00000187908 | 8 |
| GYPE | ENSG00000197465 | 8 |
| NBAT1 | ENSG00000260455 | 8 |
| TESC | ENSG00000088992 | 7 |
| PIK3C2G | ENSG00000139144 | 8 |
| CCDC198 | ENSG00000100557 | 8 |
| KRT13 | ENSG00000171401 | 8 |
| GBP6 | ENSG00000183347 | 8 |
| AQP3 | ENSG00000165272 | 8 |
| CYP2C18 | ENSG00000108242 | 8 |
| GDA | ENSG00000119125 | 8 |
| SERPINB2 | ENSG00000197632 | 8 |
| KRT4 | ENSG00000170477 | 8 |
| MSLN | ENSG00000102854 | 8 |
| SCARA5 | ENSG00000168079 | 8 |
| OSR1 | ENSG00000143867 | 8 |
| FOXI1 | ENSG00000168269 | 8 |
| ASCL2 | ENSG00000183734 | 8 |
| BSND | ENSG00000162399 | 8 |
| POU2F3 | ENSG00000137709 | 8 |
| GNG13 | ENSG00000127588 | 8 |
| GNAT3 | ENSG00000214415 | 8 |
| AXIN2 | ENSG00000168646 | 8 |
| RSPO3 | ENSG00000146374 | 8 |
| RSPO4 | ENSG00000101282 | 8 |
| LGR4 | ENSG00000205213 | 8 |
| LGR5 | ENSG00000139292 | 8 |
| LGR6 | ENSG00000133067 | 8 |
| WNT2 | ENSG00000105989 | 8 |
| WNT2B | ENSG00000134245 | 8 |
| WNT3A | ENSG00000154342 | 8 |
| WNT4 | ENSG00000162552 | 8 |
| WNT5A | ENSG00000114251 | 8 |
| WNT5B | ENSG00000111186 | 8 |
| WNT6 | ENSG00000115596 | 7 |
| WNT7A | ENSG00000154764 | 8 |
| WNT7B | ENSG00000188064 | 8 |
| WNT9A | ENSG00000143816 | 8 |
| WNT10A | ENSG00000135925 | 8 |
| WNT11 | ENSG00000085741 | 8 |
| ROR1 | ENSG00000185483 | 8 |
| FZD4 | ENSG00000174804 | 8 |
| FZD5 | ENSG00000163251 | 8 |
| FZD6 | ENSG00000164930 | 8 |
| WIF1 | ENSG00000156076 | 8 |
| NOTUM | ENSG00000185269 | 8 |
| FGF1 | ENSG00000113578 | 8 |
| FGF2 | ENSG00000138685 | 8 |
| FGF7 | ENSG00000140285 | 8 |
| FGF9 | ENSG00000102678 | 8 |
| FGF10 | ENSG00000070193 | 8 |
| FGF11 | ENSG00000161958 | 8 |
| FGF12 | ENSG00000114279 | 8 |
| FGF13 | ENSG00000129682 | 8 |
| FGF20 | ENSG00000078579 | 8 |
| SPRY1 | ENSG00000164056 | 8 |
| SPRY2 | ENSG00000136158 | 8 |
| SPRY4 | ENSG00000187678 | 8 |
| FGFR1 | ENSG00000077782 | 8 |
| FGFR2 | ENSG00000066468 | 8 |
| FGFR3 | ENSG00000068078 | 8 |
| FGFR4 | ENSG00000160867 | 8 |
| BMP1 | ENSG00000168487 | 8 |
| BMP2 | ENSG00000125845 | 8 |
| BMP3 | ENSG00000152785 | 8 |
| BMP4 | ENSG00000125378 | 8 |
| BMP5 | ENSG00000112175 | 8 |
| BMP6 | ENSG00000153162 | 8 |
| BMP7 | ENSG00000101144 | 8 |
| INHBA | ENSG00000122641 | 8 |
| ID1 | ENSG00000125968 | 7 |
| ID2 | ENSG00000115738 | 8 |
| ID3 | ENSG00000117318 | 8 |
| ID4 | ENSG00000172201 | 8 |
| TGFB1 | ENSG00000105329 | 8 |
| TGFB2 | ENSG00000092969 | 8 |
| TGFB3 | ENSG00000119699 | 8 |
| TGIF1 | ENSG00000177426 | 8 |
| TGIF2 | ENSG00000118707 | 8 |
| LTBP1 | ENSG00000049323 | 8 |
| LTBP2 | ENSG00000119681 | 8 |
| LTBP3 | ENSG00000168056 | 8 |
| LTBP4 | ENSG00000090006 | 8 |
| IHH | ENSG00000163501 | 8 |
| DHH | ENSG00000139549 | 8 |
| SHH | ENSG00000164690 | 8 |
| HHIP | ENSG00000164161 | 8 |
| GLI2 | ENSG00000074047 | 8 |
| GLI3 | ENSG00000106571 | 8 |
| GDF10 | ENSG00000266524 | 8 |
| GDF11 | ENSG00000135414 | 8 |
| CCN2 | ENSG00000118523 | 8 |
| ANKRD1 | ENSG00000148677 | 8 |
| EGF | ENSG00000138798 | 8 |
| NRG1 | ENSG00000157168 | 8 |
| NRG2 | ENSG00000158458 | 8 |
| NRG3 | ENSG00000185737 | 8 |
| NRG4 | ENSG00000169752 | 8 |
| TGFA | ENSG00000163235 | 8 |
| EPGN | ENSG00000182585 | 8 |
| EREG | ENSG00000124882 | 8 |
| HBEGF | ENSG00000113070 | 8 |
| AREG | ENSG00000109321 | 8 |
| HGF | ENSG00000019991 | 8 |
| BTC | ENSG00000174808 | 8 |
| HES1 | ENSG00000114315 | 8 |
| HES4 | ENSG00000188290 | 2 |
| NOTCH1 | ENSG00000148400 | 8 |
| NOTCH2 | ENSG00000134250 | 8 |
| NOTCH3 | ENSG00000074181 | 8 |
| NOTCH4 | ENSG00000204301 | 8 |
| JAG1 | ENSG00000101384 | 8 |
| JAG2 | ENSG00000184916 | 8 |
| DLL1 | ENSG00000198719 | 8 |
| DLL4 | ENSG00000128917 | 8 |
| CSF1 | ENSG00000184371 | 8 |
| CSF2 | ENSG00000164400 | 8 |
| CSF3 | ENSG00000108342 | 8 |
| IL33 | ENSG00000137033 | 8 |
| IL1B | ENSG00000125538 | 8 |
| IL1A | ENSG00000115008 | 8 |
| TNF | ENSG00000232810 | 8 |
| IL6 | ENSG00000136244 | 8 |
| C3 | ENSG00000125730 | 8 |
| ACE2 | ENSG00000130234 | 8 |
| TMPRSS2 | ENSG00000184012 | 8 |
| CCL5 | ENSG00000271503 | 8 |
| CCL18 | ENSG00000275385 | 8 |
| CCL14 | ENSG00000276409 | 8 |
| IL16 | ENSG00000172349 | 8 |
| PROS1 | ENSG00000184500 | 8 |
| LIF | ENSG00000128342 | 8 |
| MIF | ENSG00000240972 | 3 |
| MDK | ENSG00000110492 | 7 |
| KITLG | ENSG00000049130 | 8 |
| VEGFA | ENSG00000112715 | 8 |
| AGRN | ENSG00000188157 | 8 |
| PDGFA | ENSG00000197461 | 8 |
| PDGFB | ENSG00000100311 | 8 |
| KDR | ENSG00000128052 | 8 |
| DEPTOR | ENSG00000155792 | 8 |
| SEMA6D | ENSG00000137872 | 8 |
| CXCL2 | ENSG00000081041 | 8 |
| ANGPT1 | ENSG00000154188 | 8 |
| EDN1 | ENSG00000078401 | 8 |
| ANGPTL1 | ENSG00000116194 | 8 |
| TUB | ENSG00000166402 | 8 |
| MERTK | ENSG00000153208 | 8 |
| TNFSF12 | ENSG00000239697 | 8 |
| RARRES2 | ENSG00000106538 | 8 |
| VEGFB | ENSG00000173511 | 8 |
| VEGFC | ENSG00000150630 | 8 |
| VEGFD | ENSG00000165197 | 8 |
| NAMPT | ENSG00000105835 | 6 |
| SEMA3C | ENSG00000075223 | 8 |
| SEMA3A | ENSG00000075213 | 8 |
| SEMA3B | ENSG00000012171 | 8 |
| SEMA3E | ENSG00000170381 | 8 |
| PLXNA2 | ENSG00000076356 | 8 |
| NRP1 | ENSG00000099250 | 8 |
| SLIT1 | ENSG00000187122 | 8 |
| ROBO1 | ENSG00000169855 | 8 |
| ROBO2 | ENSG00000185008 | 4 |
| DAG1 | ENSG00000173402 | 8 |
| SDC4 | ENSG00000124145 | 8 |
| NECTIN1 | ENSG00000110400 | 8 |
| NECTIN2 | ENSG00000130202 | 8 |
| NECTIN3 | ENSG00000177707 | 8 |
| FLRT2 | ENSG00000185070 | 8 |
| FLRT3 | ENSG00000125848 | 8 |
| LAMA1 | ENSG00000101680 | 8 |
| LAMA2 | ENSG00000196569 | 5 |
| LAMA3 | ENSG00000053747 | 8 |
| LAMA4 | ENSG00000112769 | 8 |
| LAMA5 | ENSG00000130702 | 8 |
| LAMB1 | ENSG00000091136 | 8 |
| LAMB2 | ENSG00000172037 | 8 |
| LAMB3 | ENSG00000196878 | 8 |
| LAMC1 | ENSG00000135862 | 8 |
| LAMC2 | ENSG00000058085 | 8 |
| ELN | ENSG00000049540 | 7 |
| NPNT | ENSG00000168743 | 8 |
| OGN | ENSG00000106809 | 8 |
| DCN | ENSG00000011465 | 5 |
| LUM | ENSG00000139329 | 3 |
| SPARCL1 | ENSG00000152583 | 6 |
| FNDC1 | ENSG00000164694 | 8 |
| FBLN1 | ENSG00000077942 | 8 |
| GPC3 | ENSG00000147257 | 8 |
| COL21A1 | ENSG00000124749 | 8 |
| COL16A1 | ENSG00000084636 | 8 |
| COL23A1 | ENSG00000050767 | 8 |
| COL4A1 | ENSG00000187498 | 7 |
| COL4A2 | ENSG00000134871 | 8 |
| COL4A3 | ENSG00000169031 | 8 |
| COL4A4 | ENSG00000081052 | 8 |
| COL4A5 | ENSG00000188153 | 8 |
| COL4A6 | ENSG00000197565 | 8 |
| COL6A1 | ENSG00000142156 | 8 |
| COL6A2 | ENSG00000142173 | 8 |
| COL9A1 | ENSG00000112280 | 8 |
| COL18A1 | ENSG00000182871 | 8 |
| CTHRC1 | ENSG00000164932 | 8 |
| NID1 | ENSG00000116962 | 8 |
| COL6A3 | ENSG00000163359 | 8 |
| COL5A2 | ENSG00000204262 | 8 |
| COL24A1 | ENSG00000171502 | 8 |
| COL12A1 | ENSG00000111799 | 8 |
| COL28A1 | ENSG00000215018 | 8 |
| ITGA4 | ENSG00000115232 | 8 |
| ITGA3 | ENSG00000005884 | 8 |
| ITGB6 | ENSG00000115221 | 8 |
| ITGA2 | ENSG00000164171 | 8 |
| ITGA8 | ENSG00000077943 | 8 |
| ITGAX | ENSG00000140678 | 8 |
| ITGA1 | ENSG00000213949 | 8 |
| ITGB4 | ENSG00000132470 | 8 |
| TOP2A | ENSG00000131747 | 8 |
| MKI67 | ENSG00000148773 | 8 |
| PCNA | ENSG00000132646 | 8 |
| FOSL1 | ENSG00000175592 | 8 |
| FOSB | ENSG00000125740 | 8 |
| JUNB | ENSG00000171223 | 5 |
| EGR1 | ENSG00000120738 | 8 |
| STAT3 | ENSG00000168610 | 8 |
| ELF3 | ENSG00000163435 | 8 |
| CREB3L2 | ENSG00000182158 | 8 |
| CEBPD | ENSG00000221869 | 6 |
| EHF | ENSG00000135373 | 8 |
| GRHL2 | ENSG00000083307 | 8 |
| ETS2 | ENSG00000157557 | 8 |
| CLDN4 | ENSG00000189143 | 8 |
| KRT8 | ENSG00000170421 | 8 |
| KRT18 | ENSG00000111057 | 8 |
| KRT7 | ENSG00000135480 | 8 |
| CALR | ENSG00000179218 | 8 |
| MYLK | ENSG00000065534 | 8 |
| POSTN | ENSG00000133110 | 8 |
| CXCL12 | ENSG00000107562 | 8 |
| CCN1 | ENSG00000142871 | 8 |
| CLCF1 | ENSG00000175505 | 8 |
| ICAM1 | ENSG00000090339 | 8 |
| PTGS2 | ENSG00000073756 | 8 |
| EPHB2 | ENSG00000133216 | 8 |
| CD44 | ENSG00000026508 | 8 |
| SERPINA1 | ENSG00000197249 | 8 |
| CD36 | ENSG00000135218 | 8 |
| PLCXD1 | ENSG00000182378 | 8 |
| SPP1 | ENSG00000118785 | 8 |
| MMP9 | ENSG00000100985 | 8 |
| CTSK | ENSG00000143387 | 8 |
| CCL3 | ENSG00000277632 | 5 |
| CCL4 | ENSG00000275302 | 7 |
| CCL8 | ENSG00000108700 | 8 |
| CXCL9 | ENSG00000138755 | 8 |
| CXCL10 | ENSG00000169245 | 8 |
| KRT14 | ENSG00000186847 | 8 |
| MMP7 | ENSG00000137673 | 8 |
| CDKN2A | ENSG00000147889 | 8 |
| CDKN1A | ENSG00000124762 | 8 |
| DCLK1 | ENSG00000133083 | 8 |
| HIF1A | ENSG00000100644 | 8 |
| NUPR1 | ENSG00000176046 | 8 |
| ALK | ENSG00000171094 | 8 |
| ROS1 | ENSG00000047936 | 8 |
| NAPSA | ENSG00000131400 | 8 |
| CEACAM5 | ENSG00000105388 | 8 |
| LCN2 | ENSG00000148346 | 8 |
| DDX21 | ENSG00000165732 | 8 |
| MAP2K3 | ENSG00000034152 | 8 |
| B3GALT5 | ENSG00000183778 | 8 |
| SYBU | ENSG00000147642 | 8 |
| CD93 | ENSG00000125810 | 8 |
| TPSAB1 | ENSG00000172236 | 4 |
| HDC | ENSG00000140287 | 8 |
| SFTPA2 | ENSG00000185303 | 7 |
| ETV5 | ENSG00000244405 | 8 |
| MET | ENSG00000105976 | 8 |
| RALYL | ENSG00000184672 | 8 |
| TFF3 | ENSG00000160180 | 8 |
| LRMP | ENSG00000118308 | 8 |
| RELA | ENSG00000173039 | 8 |
| RELB | ENSG00000104856 | 8 |
| NFKBIZ | ENSG00000144802 | 8 |
| CD74 | ENSG00000019582 | 3 |
| FAP | ENSG00000078098 | 8 |
