## Supplemental Table 3 for "A spatially-resolved blueprint of the developing human lung reveals a WNT-driven niche for basal stem cells"

| **Level 1 Annotation** | **Level 2 Annotation** | **Level 3 Annotation** |
| --- | --- | --- |
| Endothelial | Large Vessels | Arterial Endothelial |
| Endothelial | Large Vessels | Lymphatic Endothelial |
| Endothelial | Small Vessels | Capillary Endothelial |
| Epithelial | Airway Epithelium | Goblet |
| Epithelial | Airway Epithelium | Secretory |
| Epithelial | Basal | LGR5+ Basal Proximal and Small Airways |
| Epithelial | Basal | LGR5+ Basal Proximal Specific |
| Epithelial | Basal | Transitional Basal |
| Epithelial | Distal Epithelium | Bud Tip Adjacent |
| Epithelial | Distal Epithelium | Bud Tip Progenitor 1 |
| Epithelial | Distal Epithelium | Bud Tip Progenitor 2 |
| Epithelial | Distal Epithelium | Proliferative Distal Epithelium |
| Epithelial | Epibasal | KRT13+ Epibasal |
| Epithelial | Epibasal | KRT4+ Epibasal |
| Epithelial | Multiciliated | Ciliated 1 |
| Epithelial | Multiciliated | Ciliated 2 |
| Epithelial | Multiciliated | Deuterosome |
| Epithelial | Neuroendocrine | GHRL+ Neuroendocrine |
| Epithelial | Neuroendocrine | GRP+ Neuroendocrine |
| Epithelial | Proliferative Basal | Proliferative Basal |
| Epithelial | Small Airway Epithelium | Lower Airway Progenitor |
| Epithelial | Sub-Mucosal Gland | Myoepithelial |
| Epithelial | Sub-Mucosal Gland | Serous |
| Immune | Lymphoid | B-Cells |
| Immune | Lymphoid | T-Cells |
| Immune | Myeloid | Basophils |
| Immune | Myeloid | Conventional DC1 |
| Immune | Myeloid | Conventional DC2 (CD1c+) |
| Immune | Myeloid | Monocytes |
| Immune | Myeloid | Neutrophils |
| Mesenchymal | Chondrocyte Lineage | Chondroblast |
| Mesenchymal | Chondrocyte Lineage | Chondrocyte |
| Mesenchymal | Chondrocyte Lineage | Chondrocyte Precursor |
| Mesenchymal | Chondrocyte Lineage | Mature Chondrocyte |
| Mesenchymal | Distal Mesenchyme | Distal RSPO2+ Mesenchyme 1 |
| Mesenchymal | Distal Mesenchyme | Distal RSPO2+ Mesenchyme 2 |
| Mesenchymal | Distal Mesenchyme | Distal RSPO2+ Mesenchyme 3 |
| Mesenchymal | Distal Mesenchyme | Distal RSPO2+ Mesenchyme 4 |
| Mesenchymal | Mesenchyme 2 | Fibroblast |
| Mesenchymal | Mural | Pericytes |
| Mesenchymal | Mural | Vascular Smooth Muscle |
| Mesenchymal | Muscle-Like | Myofibroblast |
| Mesenchymal | Muscle-Like | Smooth Muscle |
| Mesenchymal | Proliferative Mesenchyme | Proliferative Mesenchyme 1 |
| Mesenchymal | Proliferative Mesenchyme | Proliferative Mesenchyme 2 |
| Mesenchymal | Proximal Mesenchyme | Proximal PTCH2+ Mesenchyme |
| Mesenchymal | Proximal Mesenchyme | Proximal TWIST2+ Mesenchyme |
| Mesenchymal | Mesenchyme 1 | SCARA5+ Small Airway Mesenchyme |
| Mesenchymal | Mesenchyme 1 | Small Airway PTCH2+ Mesenchyme |
| Neural | Neural Crest | Neural Crest |
| Neural | Neuron | GAP43+ Neuron |
