## Supplemental Table 4 for "A spatially-resolved blueprint of the developing human lung reveals a WNT-driven niche for basal stem cells"

Supplementary Table 4. Antibody and TSA Dilutions and Primer Sequences

| **Antibody** | **Dilution** |
| --- | --- |
| Acetylated Tublin (Cell Signaling, Cat#5335**)** | 1:500 |
| KRT13 (Abcam, Cat#Ab92551) | 1:500 |
| ECAD (BD Biosciences, Cat#610181) | 1:500 |
| TP63 (R&D Systems, Cat#BAF1916) | 1:500 |
| **Probe** | **TSA dilution** |
| Hs-GRP-C1 (ACDbio RNAscope, Cat#465261) | 1:4000 |
| Hs-CHGA-C1/C3 (ACDbio RNAscope, Cat#311111) | C1 – 1:3000  C3 – 1:3000 |
| Hs-RFX6-C3 (ACDbio RNAscope, Cat#1258501) | 1:2000 |
| Hs-SCGB3A2-C2 (Cat#549951-C2) | 1:5000 |
| Hs-SCGB1A1-C3 (ACDbio RNAscope, Cat#469971) | 1:2000 |
| Hs-SFTPB-C1 (ACDbio RNAscope, Cat#544251) | 1:2000 |
| Hs-LGR6-C1 (ACDbio RNAscope, Cat#410461) | 1:2000 |
| Hs-TP63-C2 (ACDbio RNAscope, Cat#601891) | 1:4000 |

| Hs-LGR5-C3 (ACDbio RNAscope, Cat#311021) | 1:4000 |
| --- | --- |
| **Primer** | **Sequence** |
| 18S | F: GCAGAATCCACGCCAGTACAAG R: GCTTGTTGTCCAGACCATTGGC |
| AXIN2 | F: AGTGTGAGGTCCACGGAAAC R: CTGGTGCAAAGACATAGCCA |
| KRT13 | F: AGGTGAAGATCCGTGACTGG R: AGGGCCAGCTCATTCTCATA |
| TP63 | F: CCACAGTACACGAACCTGGG  R: CCGTTCTGAATCTGCTGGTCC |
| KRT5 | F: CTGGTCCAACTCCTTCTCCA R: GGAGCTCATGAACACCAAGC |
| SFTPB | F: GGGTGTGTGGGACCATGT  R: CAGCACTTTAAAGGACGGTGT |
| SCGB3A2 | F: AAGCTGGTAACTATCTTCCTGCT  R: AGGGGCACTTTGTTGATGAGG |
| SCGB1A1 | F: ATGAAACTCGCTGTCACCCT  R: GTTTCGATGACACGCTGAAA |
| MUC5AC | F: GCACCAACGACAGGAAGGATGAG R: CACGTTCCAGAGCCGGACAT |
| FOXJ1 | F: CAACTTCTGCTACTTCCGCC R: CGAGGCACTTTGATGAAGC |
| LGR5 | F: GTTTCCCGCAAGACGTAACT  R: CAGCGTCTTCACCTCCTACC |
| LGR6 | F: CAAGCCCTGGATCTTAGCTG  R: TTTTGGGAAACTGTCCTTGG |
